# Redirecting full-length FLT1 expression towards its soluble isoform promotes postischemic angiogenesis

**DOI:** 10.1101/2024.09.19.613989

**Authors:** Maja Bundalo, Sandra Vorlova, Jessica Ulrich, Ruggero Barbieri, Leon Richter, Leonie Höna, Manuel Egg, Julian Bock, Sarah Schäfer, Núria Amézaga Solé, Annabelle Rosa, Giuseppe Rizzo, Clemént Cochain, Wolfgang Kastenmüller, Erik Henke, Boris V. Skryabin, Timofey S. Rozhdestvensky, Moritz Wildgruber, Kristina Lorenz, Michaela Kuhn, Alma Zernecke

**Affiliations:** Institute for Experimental Biomedicine, University Hospital Würzburg, 97080 Würzburg, Germany; Institute of Pharmacology and Toxicology, University of Würzburg, Versbacher Str. 9, 97078 Würzburg, Germany; Würzburg Institute of Systems Immunology, Max Planck Research Group, Julius-Maximilians University of Würzburg, 97078 Würzburg, Germany; Institute of Anatomy and Cell Biology, University of Würzburg, 97070 Würzburg, Germany; Department of Medicine, Core Facility Transgenic Animal and Genetic Engineering Models (TRAM), University of Münster, 48149 Münster, Germany; Department of Radiology, University Hospital, LMU Munich, Munich, Germany; Leibniz-Institut für Analytische Wissenschaften-ISAS-e.V., Bunsen-Kirchhoff-Str. 11, 44139 Dortmund, Germany; Comprehensive Heart Failure Center, University Hospital of Würzburg, Am Schwarzenberg 15, 97078 Würzburg, Germany; Institute of Physiology, University of Würzburg, 97070 Würzburg, Germany

**Keywords:** angiogenesis, myocardial infarction, vasculogenesis, VEGF

## Abstract

Vascular endothelial growth factors and their tyrosine kinase receptors are key mediators of vasculogenesis and angiogenesis with FLT1 (VEGFR1) serving as a decoy receptor. A truncated mRNA transcript encoding soluble (s) FLT1 can be generated by premature cleavage and polyadenylation (APA). Although a shortening of transcripts is described in pathological settings, including heart diseases, the functional in vivo impact of FLT1 gene isoform generation and relevance for angiogenesis remain unknown. Here, we show that specific splice site mutations within Flt1 inhibit telescripting and activate APA in vivo to efficiently modulate gene isoform expression, inducing a complete loss of full-length (fl) Flt1 and a switch towards sFlt1 in mice. FLT1 is a high-affinity decoy receptor of VEGF limiting vessel overgrowth. We show that sFLT1 was sufficient for developmental vasculogenesis, whereas flFLT1 controlled ischemia-driven angiogenesis. Our results demonstrate that telescripting is essential in vivo for controlling Flt1 isoform expression and angiogenesis and can be harnessed to improve reparative revascularization. Furthermore, given the widespread abundance of APA signals, our approach may serve as a blueprint for studying telescripting and generating other truncated gene isoforms in vivo.

## Introduction

RNA processing includes capping, splicing, cleavage, and polyadenylation at the 3’-end of the pre-mRNA transcript. However, an estimated ∼70-75% of human genes contain alternative polyadenylation (APA) signals (1). Whereas APA in the last exon generates mRNA isoforms that encode the same protein but with different 3’ untranslated regions (UTRs), activation of intronic APA leads to truncated protein isoforms with distinct 3’ UTRs. APA signals are inhibited by the small nuclear ribonucloparticle (snRNP) U1 by base-pairing to 5’ splice sites, a nuclear RNA surveillance function termed ‘telescripting’ (2, 3). Widespread alterations in the APA landscape have been demonstrated in cancer and other diseases (4–6). Changes in the 3’ UTR length have also been described in the hypertrophic and failing heart, and after myocardial infarction (MI) in mice (7–9).

Vascular endothelial growth factors and their tyrosine kinase receptors VEGFR1/FLT1 and VEGFR2/KDR are key mediators of embryonic vasculogenesis and angiogenesis (10, 11). Whereas VEGFR2 is primarily expressed in endothelial cells (EC), where it regulates cell proliferation, chemotaxis and cell survival (12), FLT1 (VEGFR1), which binds VEGF-A (VEGF) with a 10-fold higher affinity than VEGFR2, is more widely expressed, including in EC, vascular smooth muscle cells and macrophages. It mainly serves as a decoy receptor without effective signal transduction in the endothelium (13–15). Notably, a truncated but stable mRNA encoding a shortened FLT1 transcript can be produced upon activating an APA signal within intron 13 of *FLT1* mRNA (3). This naturally occurring truncated FLT1, lacking the transmembrane spanning and intracellular region, is secreted as a soluble isoform (sFLT1). Because sFLT1 retains the VEGF-binding extracellular domain, it can still bind VEGF, and interact with full-length VEGF receptors thus forming non-signalling homo- and heterodimers, thereby sequestering VEGF and reducing receptor kinase signalling (16). Mice lacking VEGFR2 die *in utero* due to severe deficiencies in vascular development (17). In contrast, mice deficient in FLT1 display excessive angioblast proliferation and disorganized vessels with increased number of endothelial cells (15). The cytoplasmic signaling function of full length (fl) FLT1 appears dispensable for its decoy function, as mice with a targeted deletion of the tyrosine kinase domain of FLT1 are viable with a nearly normal vasculature (18). flFLT1 thus primarily functions as a sink to VEGF, indirectly regulating signalling of VEGF through VEGFR2. The contribution of sFLT1 versus flFLT1 to this regulation, however, is unknown.

Notably, circulating levels of sFLT1 are significantly increased in patients with acute MI and are an independent predictor for the development of acute severe heart failure (19). The contribution of telescripting to *Flt1* gene isoform expression *in vivo* and in the context of inflammation after MI, however, is unclear.

Here, we set out to investigate 1) how telescripting regulates the balance between flFlt1 and sFlt1 isoforms *in vivo*, and 2) whether this influences pathophysiological angiogenesis. We could confirm increased sFLT1 levels in mouse serum after myocardial infarction (MI) and found a shift in the expression of transmembrane *flFlt1* towards *sFlt1* in hypoxic endothelial cells. To define the role of telescripting to this shift in *Flt1* isoform expression, we generated mice with a mutated 5’ splice site in intron 13 of *Flt1* to site-specifically impede binding of U1. This led to a complete loss of the transmembrane flFLT1 receptor and a concomitant switch towards expression of the truncated sFLT1 isoform. These data demonstrate for the first time that telescripting is a key regulator of the flFLT1/sFlt1 expression balance *in vivo*. By completely switching the endogenous expression from *flFlt1* towards *sFlt1* in our novel mouse model, we furthermore pinpoint that expression of sFLT1 only (in the absence of flFLT1) is sufficient for normal developmental vasculogenesis and postnatal angiogenesis, whereas flFLT1 constrains post-ischemic angiogenesis in the adult.

## Results

### Differential regulation of flFlt1 and sFlt1 mRNA in tissue and endothelial cells in response to myocardial injury and hypoxia

sFLT1 can be produced upon activation of an APA signal within intron 13 of *Flt1* mRNA (**Figure 1A**) (3). Absolute mRNA expression levels of both *sFlt1* and *flFlt1* isoforms were the highest in mouse hearts and kidneys (**Figure 1B**). Patients with MI have been reported to display increased circulating levels of sFLT1 (19). In line, we noted elevated sFLT1 serum concentrations in mice after experimental MI induced by the permanent ligation of the left anterior descending coronary artery (**Figure 1C**). Next, we evaluated the impact of hypoxia on *Flt1* gene isoform expression in ECs. In primary murine EC rendered hypoxic, as confirmed by the induction of VEGF-A, a significant increase in the ratio between *sFlt1* and *flFLT1* was observed (**Figure 1D**). A similar shift could be observed in 3’RNA-seq results showing increased reads at defined poly(A) sites within *Flt1* intron 13 under hypoxic conditions (**Figure 1E**). The contribution of telescripting to the regulation of these *Flt1* isoforms and their functional role in post-ischemic angiogenesis and tissue repair remains unclear.

**Figure 1.**
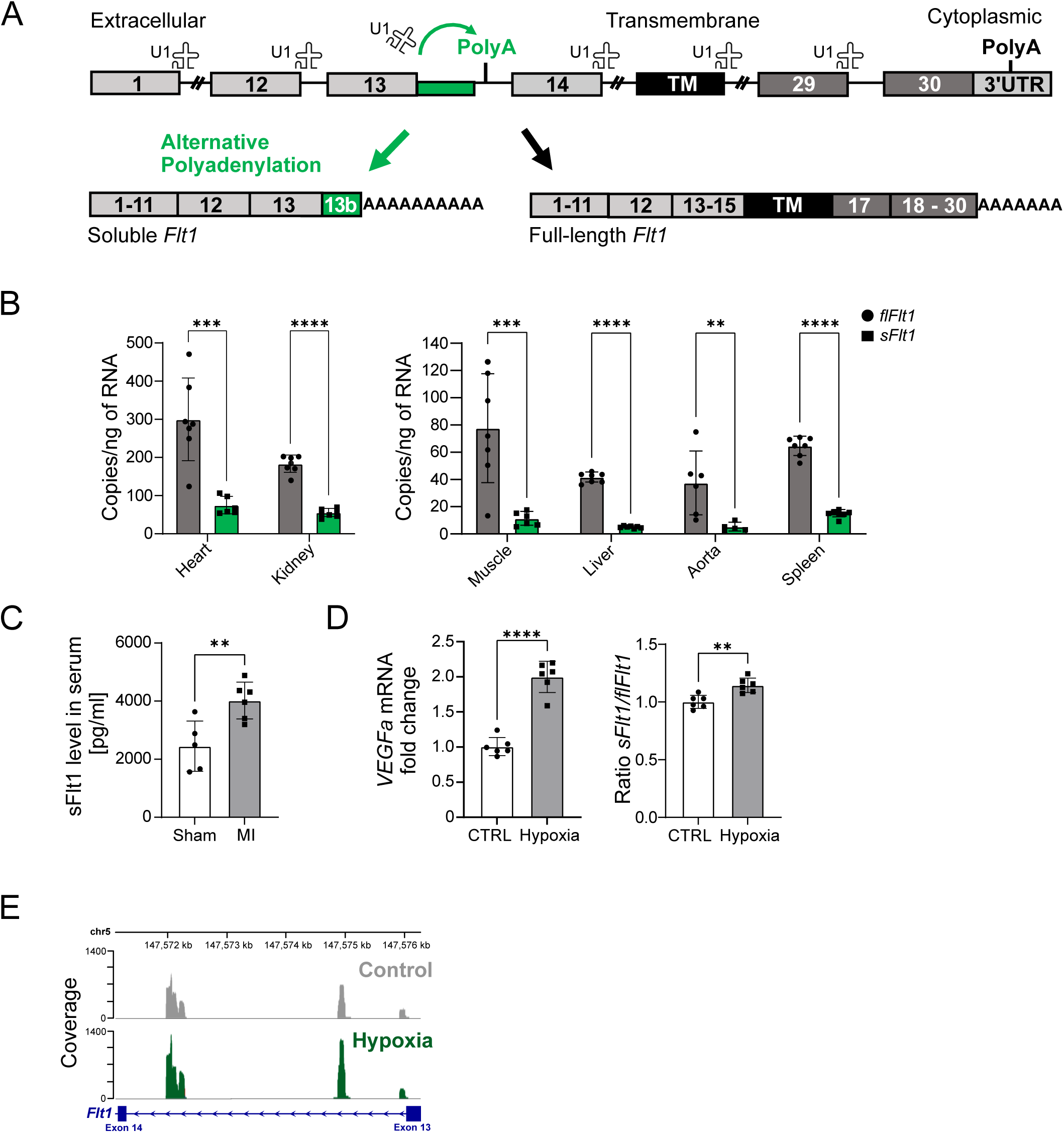
Expression of flFlt1 and sFlt1 mRNA is differentially regulated under hypoxia and in response to myocardial injury. (**A**) Scheme of *Flt1* genomic structure.Interfering with telescripting by impeding binding of U1 snRNP to the 5’ splice site in intron 13 leads to the activation of an alternative cleavage-polyadenylation (polyA) signal within intron 13, resulting in a stable truncated transcript with partial retention of intron 13. (**B**) Absolute number of mRNA copies of flFlt1 and sFlt1 in indicated Flt1^+/+^ tissues (n=5-7 mice), analyzed using gBlocks as a standard. **(C)** Serum level of *sFlt1* in serum of mice at day 1 after MI (n=5 mice) compared to sham controls (n=6 mice) and (**D**) qPCR analysis of *Vegf-a* expression and *sFlt1*/*flFlt1* ratio in lung endothelial cells exposed to hypoxia over 24 h (n=6 experiments performed with pooled endothelial cells isolated from n=8 mice). (**E**) Differential usage of *Flt1* Intron 13 polyA sites (PAS) in control and hypoxia exposed endothelial cells. Data represent mean ± SD. Results were analyzed by two-tailed, unpaired Student t test. **p<0.01 and ***p<0.001, ****p<0.0001. flFlt1-full length Flt1, sFlt1 – soluble Flt1, MI – myocardial infarction.

### Introducing selective point mutations in Flt1^In13pA^ mice activates APA forcing both selective expression of soluble Flt1 and knockdown of the full length receptor

Alternative *Flt1* pre-mRNA processing involves retention of intron 13 and activation of intronic APA signals to produce *sFlt1* (**Figure 1A**). Sterically blocking binding of the splicing factor U1 to the *Flt1* exon13/intron13 junction, by a complementary morpholino oligonucleotide (MoVE1), dose-dependently induced activation of the alternative polyA site in intron 13 and thus the generation of *sFlt1* in a mouse endothelial cell line (SVEC) *in vitro* (**Supplementary Figure 1A, B**), similar to previous findings in human cells (3). Intron 13 nucleotide sequence of *Flt1* gene is highly conserved between species, resulting in a highly conserved C-terminus of sFLT1 protein (**Supplementary Figure 1C, D**). We hypothesized that disrupting the binding of U1 to the *Flt1* exon13/intron13 junction by specific point mutations could thus site-specifically interfere with telescripting, leading to a shift in the balance between *sFlt1* and *flFlt1* expression levels. To test this, we created a mouse model by mutating the *Flt1* exon13/intron13 5’ splice site by homology-directed repair using CRISPR/Cas9 gene editing with the intent to disrupt the U1 consensus binding sequence (**Figure 2A**). Introduced silent mutations in exon 13 (−4C and −2C) and intron 13 (+2C and +5A) were chosen to maintain the FLT1 amino acid sequence during translation.

**Figure 2.**
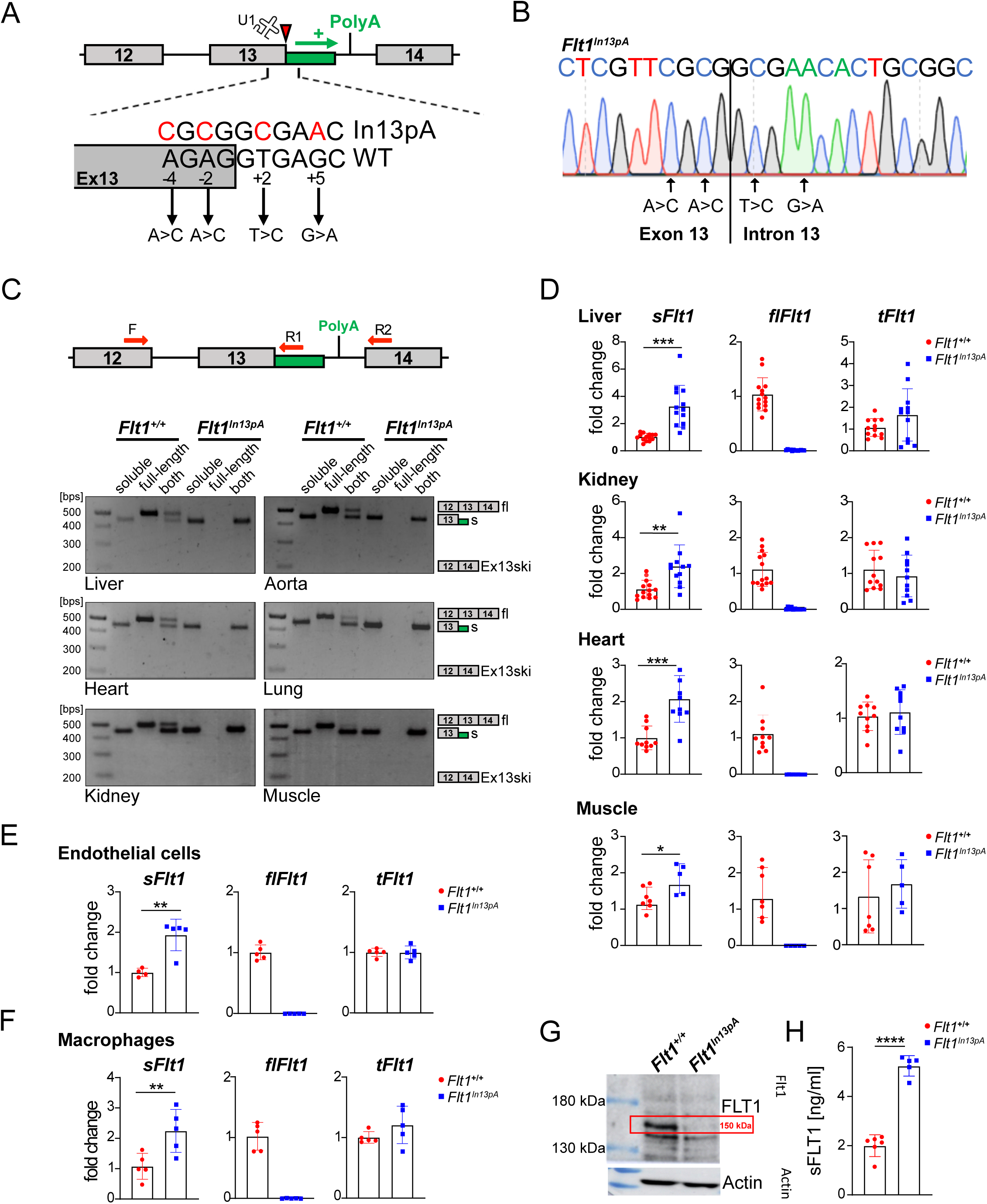
Homozygous *Flt1^In13pA^* mice lack *flFlt1* and display a complete switch towards *sFlt1* expression. **(A)** Introduction of silent genomic mutations in Flt1 exon 13 (−4C and −2C) and in intron 13 (+2C and +5A) by CRISPR/Cas9, **(B)** as confirmed by sequencing of genomic DNA from homozygous Flt1^In13pA/In13pA^ mice. (**C**) PCR products amplified with primers specific for soluble *Flt1* (F+R1), full length *Flt1* (F+R2), and both *Flt1* isoforms (F+R1+R2); PCR products using cDNA templates isolated from different organs were analyzed by gel electrophoresis. Ex13SKI, exon 13 skipping. (**D-F**) qPCR quantification of *sFlt1*, *flFlt1*, and *tFlt1* transcript expression in **(D)** different tissues (n=7-14 *Flt1^+/+^* and n=5-13 *Flt1^In13pA^* mice), **(E)** isolated lung endothelial cells (independent endothelial cell isolations from n=4 *Flt1^+/+^* and n=5 *Flt1^In13pA^* mice) and **(F)** bone marrow-derived macrophages (independent BMDM cultures from n=5 mice per genotype). (**G-H**) Untreated *Flt1^+/+^* and *Flt1^In13pA^* lung endothelial cells were analyzed for **(G)** FLT1 expression in cell lysates by Western blotting (pooled endothelial cells isolated from n=8 mice per genotype, one representative of 2 independently performed experiments), and for **(H)** sFLT1 levels in cell culture supernatant by ELISA (n=5 *Flt1^+/+^* and n=6 *Flt1^In13pA^* mice). Data represent mean ± SD. Results were analyzed by two-tailed, unpaired Student t-test. *p<0.05, **p<0.01, ***p<0.001, ****p<0,0001.

F1 founder mice heterozygous for the *Flt1*^In13pA^-allele with all four point mutations were genotyped by TaqMan Assays (Supplementary Figure 2A and B) and subsequently bred to homozygosity (**Figure 2B**). Crossing of heterozygous mice gave rise to offspring close to the expected mendelian ratio of 1:2:1 (*Flt1*^+/+^: 67, *Flt1*^+/In13pA^: 134, *Flt1*^In13pA/In13pA^: 78, **Supplementary Figure 2C**). To analyze if mutating the U1 binding site is sufficient to change specifically the APA-dependent balance of s*Flt1* and fl*Flt1* in *Flt1*^In13pA/In13pA^ mice (hereafter referred to as *Flt1*^In13pA^ mice), cDNA from the aorta, heart, liver, lung, kidney, and muscle was amplified by non-quantitative three-oligo PCR using two reverse primers specific for the full-length or the truncated soluble isoform respectively, together with a common forward primer. PCR products could be detected for both *Flt1* isoforms in all tissue samples in *Flt1*^+/+^ animals. In contrast, *flFlt1* was undetectable in *Flt1*^In13pA^ samples, whereas *sFlt1* transcripts were still clearly amplified (**Figure 2C**). Previous experiments had shown that steric hindrance of 5’ splice sites leads to exon skipping or cryptic splice site usage if an alternative intronic polyA signal is not present (20). Both alternative splicing events can be excluded since neither an exon 13 skipped product nor any other alternative spliced products could be detected using primers surrounding exon 13 for amplification (**Figure 2C**).

To precisely characterize the impact of the introduced 5’ splice site mutations on *Flt1* pre-mRNA processing and subsequent protein expression, we performed 3’RACE on RNA isolated from *Flt1*^In13pA^ liver tissue. Sequencing of the oligo(dT) primed product revealed a read-through of 115 nucleotides in intron 13 followed by cleavage and polyadenylation (**Supplementary Figure 2D**). Of these, 96 nucleotides encode a unique 32 amino acid sFLT1 tail (including the stop codon). Of note, besides truncation of the protein-coding sequence and addition of a novel tail, the introduced 5’ splice site mutation enforcing APA in the *Flt1* gene also changed the 3’ UTR of the resulting mRNA, which may influence the stability, localization, export and translational efficiency of the transcript.

We thus analyzed tissue samples by real-time RT-qPCR to quantify changes in *Flt1* isoform balance. Tissue from *Flt1*^In13pA^ mice showed a significant (approximately 2-fold) increase in *sFlt1* mRNA expression compared to *Flt1^+/+^* mice, whereas transcripts for *flFlt1* were again undetectable. Expression of total *Flt1* transcripts was unaltered, supporting a switch in isoform expression (**Figure 2D**). Similar gene expression patterns were found in ECs isolated from the lung, as well as in bone marrow-derived macrophages (BMDM) from *Flt1*^In13pA^ versus control animals (**Figure 2E, F**). Western blot analyses of EC cell lysates furthermore confirmed complete loss of transmembrane flFLT1 protein in *Flt1*^In13pA^ ECs (**Figure 2G**). *Flt1*^In13pA^ EC cultivated for 28h showed a 2.5-fold increase of sFLT1 protein release in the culture supernatant compared to control as measured by ELISA (**Figure 2H**). These analyses show that genomic mutations blocking U1 snRNP binding to a specific 5’splice site are sufficient to induce a shift in pre-mRNA processing of *Flt1* by site-specifically interfering with telescripting, triggering a complete switch in the expression of the full-length receptor to its endogenous soluble decoy isoform.

### Flt1*^In13pA^* mice exhibit normal vascular development

Whereas knockout of *Kdr* leads to embryonic lethality due to a lack of vasculogenesis (17), loss of *Flt1* as a negative regulator of angiogenesis results in embryonic death due to an overshooting growth of blood vessel ECs (15). As sFLT1 can still sequester VEGF to limit VEGFR2 signaling (13, 14), we expected a non-lethal phenotype, with potentially mild defects in vessel formation in our mice selectively expressing sFLT1 and lacking flFLT1. However, we did not detect any obvious defects during embryonic development, and embryo morphology and crown-rump length were unaltered (**Figure 3A, B**). Similarly, postnatal retinal vascular development, evaluated using flat-mounted retinas after fluorescein isothiocyanate (FITC)-conjugated isolectin-B4 staining, was unaltered, with no significant differences in the vascular area, number of branch points or the total vessel length in *Flt1*^In13pA^ as compared to *Flt1*^+/+^ mice at 5 and 7 days *post partum* (**Figure 3C**). Moreover, weight relative to body weight (**Figure 3D**) and morphology of heart, liver, and kidney was unaltered in 8-week-old mice (**Supplementary Figure 3A**). No abnormalities or differences in the number of CD31^+^ vessels could be detected in histological sections in these organs of adult *Flt1*^In13pA^ mice compared to controls (**Figure 3E, F**, and **Supplementary Figure 3B**). These data provide unprecedented and unequivocal evidence that the presence of sFLT1 in the absence of flFLT1 is sufficient for normal developmental vasculogenesis and pre- and postnatal angiogenesis.

**Figure 3.**
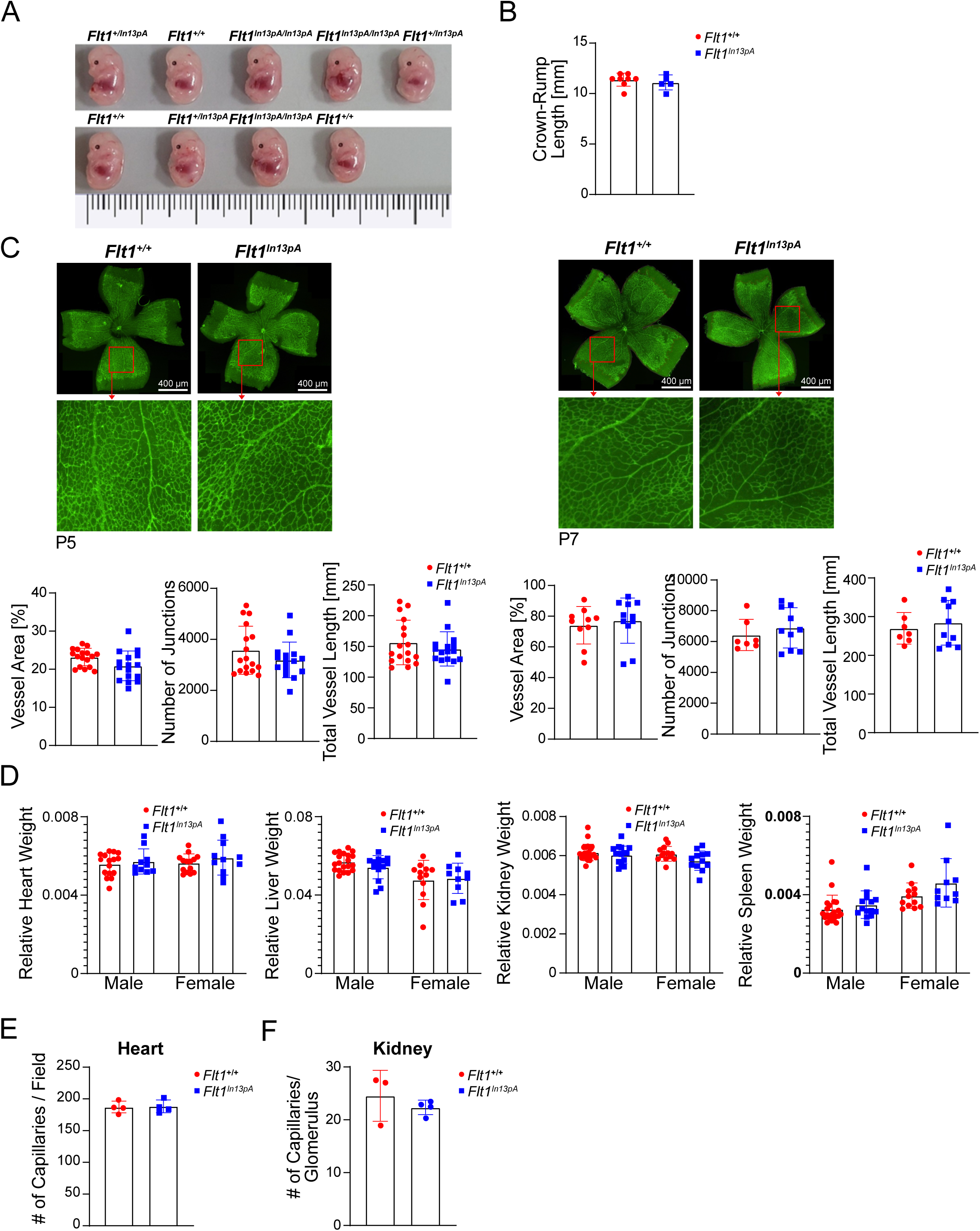
Embryo size and developmental angiogenesis are unaffected in *Flt1^In13pA^* mice. (**A**) Representative photographs of littermate embryos from heterozygous (*Flt1^+/In13pA^*) mouse breeding, and (**B**) crown rump length of *Flt1^+/+^* and homozygous *Flt1^In13pA^* embryos at E13.5 (n=8 *Flt1^+/+^* and n=5 *Flt1^In13pA^* mice). (**C**) Analysis of vessel area, junction numbers, and total vessel length of isolectin B4-stained retinal blood vessels at day 5 (P5, n=16 *Flt1^+/+^* and n=17 *Flt1^In13pA^* mice) and day 7 (P7, n=7-10 *Flt1^+/+^* and n=10-11 *Flt1^In13pA^* mice). (**D**) Weight of heart, liver, kidney and spleen relative to total body weight in male and female 8-week-old *Flt1^+/+^* and *Flt1^In13pA^* mice (n=10-23 Flt1^+/+^ and n=10-15 Flt1^In13pA^ mice). (**E, F**) Number of CD31 positive capillaries per microscopic field in **(E)** tissue sections through the heart (n=4 mice per genotype) and **(F)** kidney glomeruli (n=3 *Flt1^+/+^* and n=4 *Flt1^In13pA^* mice) of 8-week-old male *Flt1^+/+^* and *Flt1^In13pA^* mice. Data represent mean ± SD. Results were analyzed by two-tailed, unpaired Student t test.

### Redirection of flFlt1 towards sFlt1 isoform expression unleashes angiogenesis after injury

Different from vessel formation during development, the postnatal vascular system adapts to pathological cues by angiogenesis to maintain tissue homeostasis, i.e., by sprouting of capillaries from the pre-existing vasculature, e.g., in ischemic tissue (21). To explore whether a shift in FLT1 isoform expression may regulate pathological angiogenesis, such as after ischemic tissue injury *in vivo*, we induced hind-limb ischemia by femoral artery ligation (22) in *Flt1*^In13pA^ and *Flt1*^+/+^ mice. Notably, an increased CD31^+^ capillary density relative to the number of myocytes was evidenced at day 14 after injury in ischemic (**Figure 4A**) but not the non-ligated (**Supplementary Figure 4A**) gastrocnemius muscles of *Flt1*^In13pA^ versus *Flt1*^+/+^ mice. We also noted a reduction in macrophage accumulation in the injured muscles in *Flt1*^In13pA^ mice after injury (**Figure 4B**).

**Figure 4.**
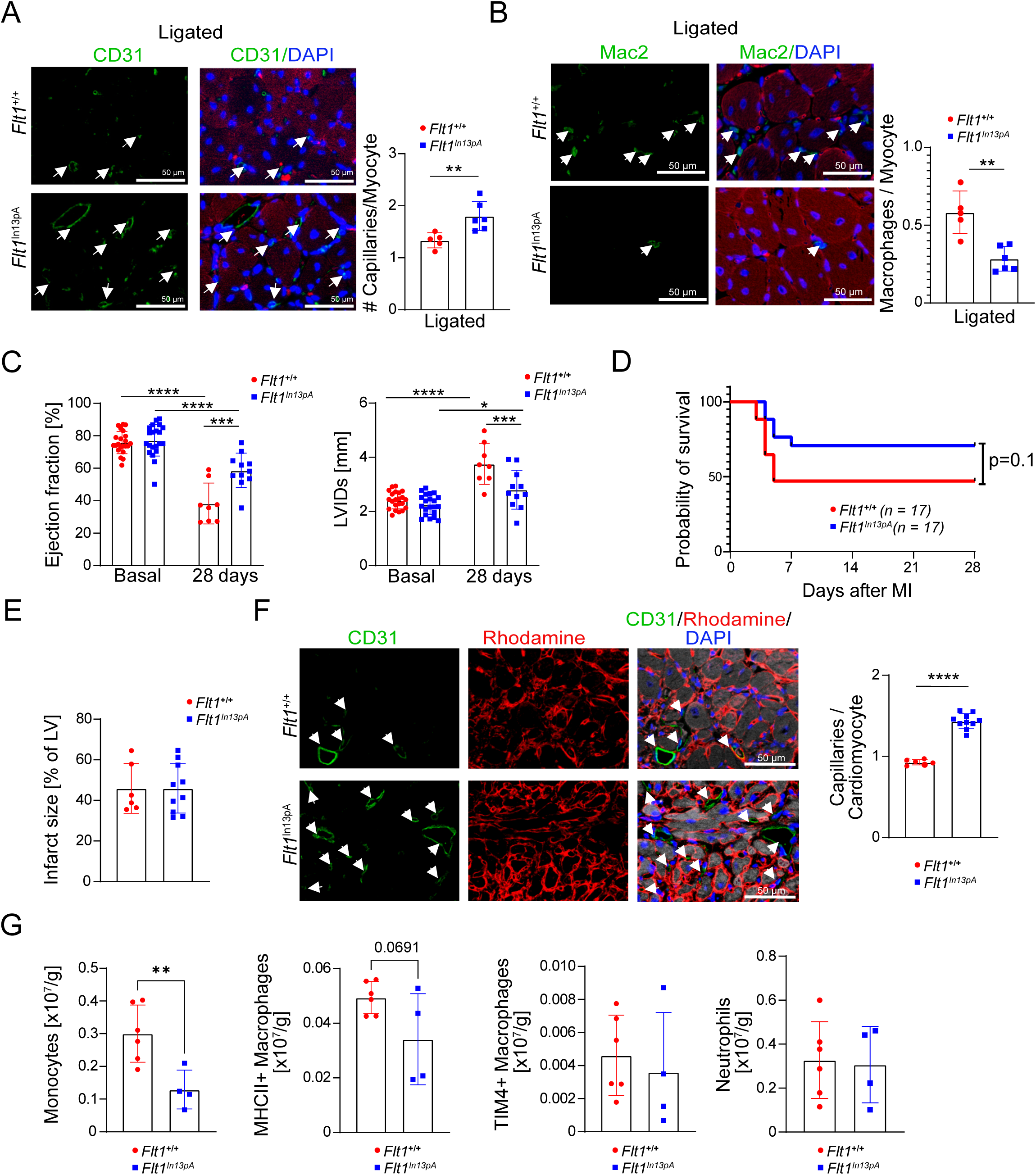
Lack of flFLT1 promotes postischemic angiogenesis and improves cardiac function after myocardial infarction. (**A, B**) Representative immunofluorescence staining and quantification of (A) CD31^+^ capillaries in ligated gastrocnemius muscle and (B) Mac2^+^ monocytes/macrophages in the ischemic gastrocnemius muscle of *Flt1*^+/+^ (n=5) and *Flt1*^In13pA^ (n=6) mice at 14 days after injury in the hindlimb ischemia model (scale bars, 50 µm). Results were analyzed by two-tailed, unpaired Student’s t-test. (**C-G**) *Flt1*^+/+^ and *Flt1*^In13pA^ mice were subjected to MI injury. **(C)** Quantification of ejection fraction (%) and internal left ventricular end-systolic diameter (mm) under basal conditions (n=22 *Flt1*^+/+^ and n=21 *Flt1*^In13pA^ mice) and at day 28 (n=8 *Flt1*^+/+^ and n=11 *Flt1*^In13pA^ mice) after MI. Data were analyzed by One way ANOVA followed by Tukey’s post-hoc test. **(D)** Kaplan-Meier survival analysis over 28 days post-MI (n=17 mice per genotype). **(E)** Quantification of the relative infarct size at 28 days post-MI (n=6 *Flt1*^+/+^ and n=10 *Flt1*^In13pA^ mice). Results were analyzed by two-tailed, unpaired Student’s t-test. **(F)** Representative immunofluorescence staining of CD31^+^ capillaries and Rhodamine staining to demarcate cardiomyocytes in the infarcted myocardium of *Flt1*^+/+^ (n=6) and *Flt1*^In13pA^ (n=10) mice at 28 days post-MI (scale bars, 50 µm). Quantification of CD31^+^ capillaries relative to the number of cardiomyocytes. Results were analyzed by two-tailed, unpaired Student’s t-test. **(G)** Quantification of the number of monocytes, MHCII^+^, TIM4^+^ macrophages and neutrophils in the infarcted myocardium at day 3 post-MI (n=6 *Flt1*^+/+^ and n=4 *Flt1*^In13pA^ mice), as assessed by flow cytometry. Results were analyzed by unpaired Student t test. Data represent mean ± SD. **p<0.01, ***p<0.001, ****p<0.0001.

We further scrutinized the angiogenic response after ischemic injury in mice subjected to permanent MI. Left ventricular function, assessed by high-resolution echocardiography, was similar between groups at baseline. After MI, Flt1In13pA mice were protected from the pronounced contractile dysfunction observed in Flt1+/+ mice, as evident from a stronger ejection fraction (EF) (**Figure 4C**) and fractional shortening (FS) (**Supplementary Figure 4C**) and a smaller left ventricular inner diameter in systole (LVID) (**Figure 4C**). Heart rates were comparable (**Supplementary Fig. 4D**). Kaplan–Meier analysis showed a trend towards improved survival over 28 days post-MI (**Figure 4D**). No differences were noted in infarct size between genotypes (**Figure 4E**). Notably, the capillary density in the infarct border zone was increased in *Flt1*^In13pA^ mice (**Figure 4F**) but not in the remote area (**Supplementary Figure 4E**). Flow cytometric analysis of the cardiac inflammatory cell infiltrate at day 3 after MI furthermore demonstrated a reduction in monocyte accumulation and a trend towards diminished MHCII^+^ macrophage counts but unaltered neutrophil and resident TIM4^+^ macrophage levels (**Figure 4G**, and **Supplementary Figure 4F**). These data indicate that a shift in the expression from flFLT1 towards sFLT1 lowers monocyte recruitment while accelerating angiogenesis and tissue repair after ischemic injury.

### flFLT1 constrains angiogenesis in vitro

To further mechanistically dissect the impact of the ratio between soluble and full-length FLT1 on angiogenesis, we used primary ECs isolated from the lungs of *Flt1*^In13pA^ and *Flt1*^+/+^ mice in a tube formation assay (**Supplementary Figure 5A**). In line with the enhanced angiogenic response observed *in vivo*, an increased number of tubes formed in *Flt1*^In13pA^ compared to *Flt1*^+/+^ EC cultures (**Figure 5A**), and Western blot analyses showed an increased VEGFR2 phosphorylation in *Flt1*^In13pA^ compared to *Flt1*^+/+^ ECs, without differences in *Kdr* mRNA or total VEGFR2 protein levels (**Figure 5B**, data not shown). Bulk RNA sequencing and gene set enrichment analyses furthermore unveiled a significant upregulation in genes controlling angiogenesis (annotated in the databases Molecular Signatures Database, MSigDB, and Gene Set Enrichment Analysis, GSEA) and an increased expression of e.g. C-X-C chemokine receptor type 4 (*Cxcr4*), nitric oxide synthase 3 (*Nos3*), delta-like 4 (*Dll4),* and angiopoietin 2 (*Angpt2*) (23) in hypoxic *Flt1*^In13pA^ compared to *Flt1*^+/+^ EC (**Supplementary Figure 5B, C**). Cell proliferation was increased in *Flt1*^In13pA^ compared to *Flt1*^+/+^ ECs (**Supplementary Figure 5D**). These findings were obtained despite increased levels of sFLT1 in the supernatants of *Flt1*^In13pA^ compared to *Flt1*^+/+^ EC cultures (**Figure 2H**). They are in opposition to previous findings that addition of sFLT1 to EC culture medium represses tube formation (24). To further address this issue, we next analyzed the VEGF-dependent angiogenic effects in *Flt1*^+/+^ ECs grown in *Flt1*^+/+^ EC-conditioned medium (containing sFLT1 at a concentration of 2000 pg/ml) and *Flt1*^In13pA^ ECs in *Flt1*^In13pA^ EC-conditioned medium (sFLT1 at 5242 pg/ml). In response to exogenous VEGF, *Flt1*^In13pA^ cells again formed more tubes than *Flt1*^+/+^ cells, even though exposed to conditioned medium containing higher amounts of sFLT1 (**Supplementary Figure 5E**). In contrast, when investigating the VEGF-dependent angiogenic effect in SVEC cells grown in either supernatant from *Flt1*^+/+^ or *Flt1*^In13pA^ cells, we observed the expected inhibition of tube formation in the presence of *Flt1*^In13pA^ EC-conditioned medium as compared to responses elicited in *Flt1*^+/+^ EC-conditioned medium (**Figure 5C**). These data together indicate that *Flt1*^In13pA^ ECs have a higher intrinsic angiogenic potential, presumably linked to the lack of suppressive flFLT1 signaling in ECs, despite the increased expression of sFLT1. This finding highlights the functional relevance of the control of Flt1 isoform levels regulated by telescripting. Interestingly, a recent report predicted high synthesis rates of sFLT1 in ECs and in tissue (25), substantiating the notion that the cell-intrinsic balance of full-length and soluble FLT1 fine-tunes EC responses to VEGF.

**Figure 5.**
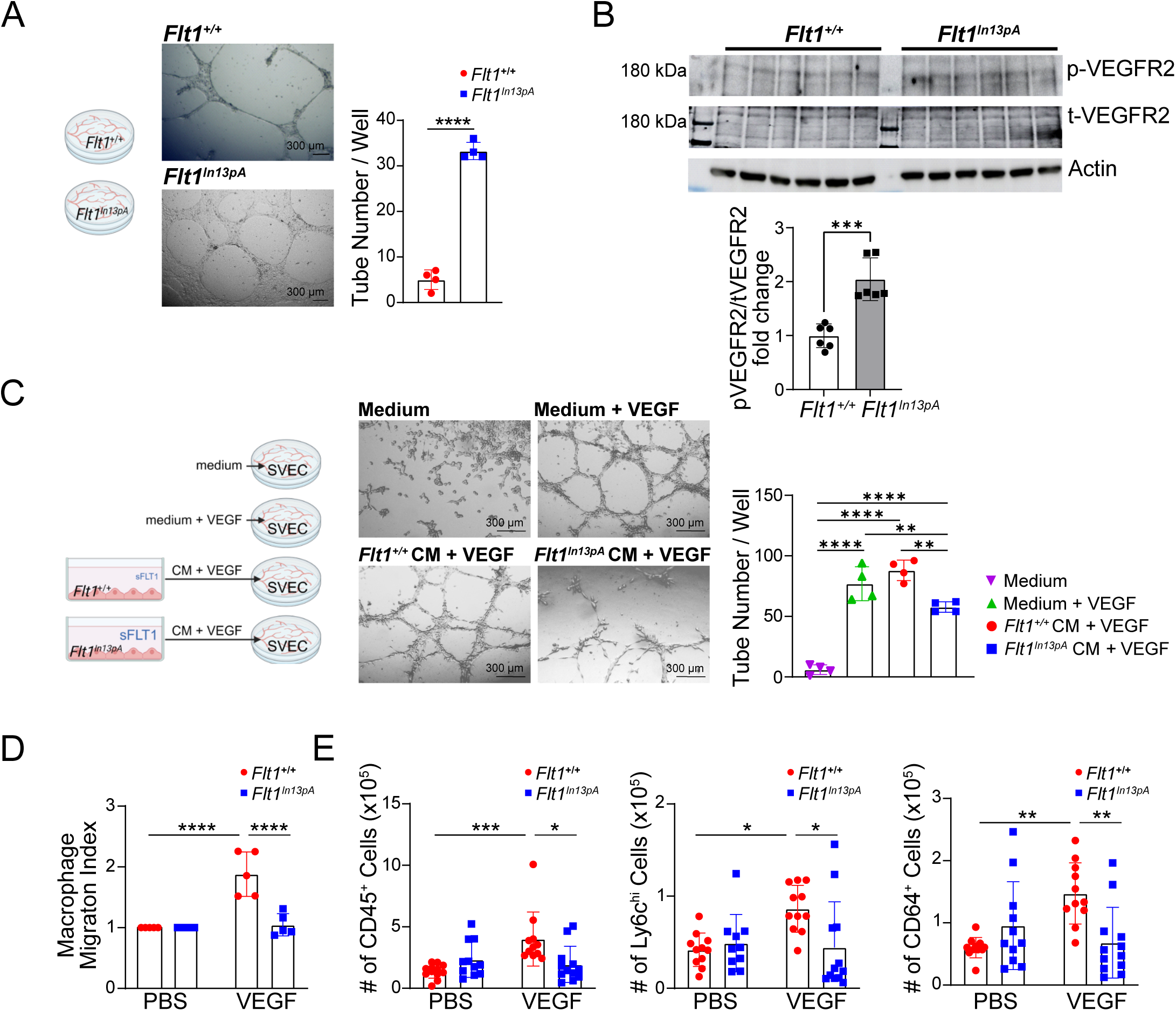
flFLT1 controls endothelial cell proliferation and monocyte/macrophage migration. (**A**) Tube formation assay of *Flt1^+/+^* and *Flt1^In13pA^* mouse lung endothelial cells cultured for 24h (independent endothelial cell isolations from n=4 mice per genotype). Representative phase contrast micrographs (scale bars, 300 µm), and quantification of tubes. Results were analyzed by two-tailed, unpaired Student’s t test. (**B**) Phosphorylated VEGFR2 (p-VEGFR2) and total VEGFR2 (t-VEGFR2) was analyzed in untreated *Flt1^+/+^* and *Flt1^In13pA^* mouse lung endothelial cell lysates by Western blotting (n=6 per genotype). Results were analyzed by two-tailed, unpaired Student’s t test. (**C**) Tube formation assay of SVEC incubated with unconditioned media, VEGF-supplemented unconditioned media, or VEGF-supplemented conditioned medium (CM) of *Flt1^+/+^* or *Flt1^In13pA^* lung endothelial cells (n=4 per condition). Representative phase contrast micrographs, (scale bars, 300 µm), and quantification of tubes. Results were analyzed by One way ANOVA followed by Tukey’s post-hoc test. (**D**) Chemotaxis of *Flt1^+/+^* and *Flt1^In13pA^* bone marrow-derived macrophages towards VEGF compared to vehicle (independent BMDM cultures from n=5 mice per genotype). Results were analyzed by One way ANOVA followed by Tukey’s post-hoc test. (**E**) Number of CD45^+^ leukocytes, Ly6C^hi^ monocytes, and of CD64^+^ macrophages in the air pouch of *Flt1^+/+^* and *Flt1^In13p^* mice, injected with PBS or VEGF and analyzed by flow cytometry after 24 h (n=10-12 mice per genotype). Data were pooled from 3 independent experiments. Results were analyzed by One way ANOVA followed by Tukey’s post-hoc test. Data represent mean ± SD. *p<0.05, **p<0.01, ***p<0.001, ****p<0.0001.

### flFLT1 controls VEGF-induced chemotactic responses in macrophages in vitro

FLT1 has previously been shown to mediate VEGF-induced chemotactic responses of monocytes that lack VEGFR2 expression (26). We therefore also tested whether the FLT1 isoform switch affects bone marrow-derived macrophage (BMDM) migration. While we noted equal migratory responses of *Flt1*^In13pA^ and *Flt1*^+/+^ macrophages towards CCL2 and FCS after 3.5 hours of migration (data not shown), VEGF elicited no chemotactic activity in *Flt1*^In13pA^ macrophages compared to efficient responses in *Flt1*^+/+^ macrophages (**Figure 5D**). Similarly, the VEGF-induced *in vivo* accumulation of monocytes and macrophages in an air pouch model was inhibited in *Flt1*^In13pA^ compared to *Flt1*^+/+^ mice (**Figure 5E**). These data indicate that increased expression of sFLT1, and lack of flFLT1, in *Flt1*^In13pA^ macrophages impairs their migration towards VEGF.

## Discussion

Dysregulated RNA processing has a profound impact on health and disease, with APA emerging as one of the major players. Activation of polyadenylation signals located in introns can result in stable and shortened mRNAs and the generation of C-terminal truncated protein isoforms with often antagonistic functions (3). A shortened *sFlt1* transcript can be produced dependent on an APA signal within intron 13 (3). sFlt1 has been implicated in several diseases, including heart failure (19), preeclampsia (27), and aging (28). In our study, assessing the expression of *flFlt1* and *sFlt1*, we observed an isoform-specific shift towards *sFlt1* in the infarcted heart *in vivo* and in hypoxic ECs *in vitro*. Whether this shift in gene isoform expression relies on telescripting and can be genetically targeted to modulate and study its role *in vivo*, had not been investigated prior to this work. The non-coding snRNP U1 is part of the spliceosome, but it also inhibits the activation of cryptic or APA signals by telescripting (2). With the aim to impede telescripting, we generated a mouse model where silent mutations were introduced to the 5’ splice site in intron 13 of *Flt1* using CRISPR/Cas9. These mutations indeed resulted in activation of an alternative intronic polyA signal in the adjacent intron without provoking exon skipping, even though splice site mutations have been generally described to either induce alternative splicing or RNA degradation by nonsense-mediated decay (20). Notably, APA led to a complete knockout of *flFlt1* with a concomitant switch towards increased expression of *sFlt1*, demonstrating that telescripting is essential for the generation of flFLT1 *in vivo*. sFLT1, dominantly produced in our mice, is identical to the endogenously produced isoform found *in vivo* and increased in pathologies including heart failure (19), preeclampsia (29), or aging (28).

Although it is now widely accepted that flFLT1 primarily functions as a decoy receptor to sequester excess VEGF without effective signal transduction in the endothelium (13–15), it has been difficult to fully dissect its exact role in vessel formation. Different genetically modified mouse models have been generated, including the conditional Cre-*lox*P-mediated full body knockout of FLT1 in neonatal and adult mice (30) and the deletion of a FLT1 kinase domain resulting in the expression of inactive membrane-bound flFLT1 and sFLT1 (18), both of which show vascular normalcy at steady state. Our model is the first to show a complete endogenous shift towards sFlt1 with concomitant knockout of *flFlt1*. We demonstrate that the two isoforms of FLT1 differentially affect developmental and post-natal physiological angiogenesis and reparative pathological angiogenesis in adult mice. Expression of sFLT1, in the absence of flFLT1 in *Flt1*^In13pA^ mice, was sufficient to allow the development of a normal vasculature both during development and postnatally. *Flt1*^In13pA^ mice were viable and fertile and displayed no abnormalities in organ size and vascularization. Our findings are in line with *in vitro* models demonstrating that the expression of *sFlt1* alone is sufficient to rescue branching morphogenesis in murine *Flt1*^−/−^ embryonic stem cell-derived vessels (31, 32). In contrast, tube formation and regulation of genes promoting angiogenetic cell responses in endothelial cells *in vitro*, and angiogenesis in response to ischemia in mice was highly amenable to regulation by the different FLT1 isoforms, pointing towards a cell-intrinsic dominant negative role of flFLT1 that restrains reparative angiogenesis in adult mice. A similar mechanism of inhibiting overshooting vessel growth may operate in *Flt1* tyrosine kinase-deficient (*Flt1 tk*^−/−^) mice (18), where tyrosine kinase-deficient FLT1 can still serve as a local VEGF sink and form non-signaling FLT1 and FLT1/VEGFR2 receptor dimers to inhibit angiogenic responses in ECs. Interestingly, *Flt1 tk*^−/−^ mice harbor a reduction in microvascular density after femoral artery ligation, driven by a concomitant defect in pro-angiogenic macrophage accumulation (33). Previous work has shown that monocytes/macrophages accumulate in the infarct region and promote post-MI angiogenesis (34). However, in our *Flt1*^In13pA^ mice, the shift in expression from *flFlt1* to *sFlt1* enhanced angiogenesis in the skeletal and heart muscle in response to ischemic injury. This beneficial effect was seen despite reducing macrophage accumulation in the injured tissues in *Flt1*^In13pA^ compared to *Flt1*^+/+^ mice. This suggests that lifting the flFLT1-mediated break on VEGFR2 angiogenic signaling in ECs triggered by locally increased VEGF-A bioavailability is sufficient to promote neovascularization, independent of macrophage recruitment to the tissue. In line with this model, *Vegfb* gene transfer to enhance VEGF-B binding to FLT1 increased angiogenesis in adipose tissue, possibly due to an increased VEGF-A availability for VEGFR2 binding, increased VEGFR2 homodimer and decreased FLT1/VEGFR2 heterodimer formation. (35)

Improved neovessel formation after MI mitigates scarring and worsening of heart function (36), and we also noted reduced cardiac dysfunction after MI in *Flt1*^In13pA^ mice. These data align with recent observations in zebrafish, where following cryoinjury, *Flt1*^−/−^ mutant hearts display enhanced coronary revascularization and improved cardiac regeneration (37). A shift toward increased *sFlt1* but reduced *flFlt1* expression in hypoxic ECs *in vitro* may thus extend the concept of a pro-inflammatory environment triggering vessel sprouting in wound angiogenesis (38) by modulating cellular FLT1 availability. Unlike VEGF, which provides strong angiogenic signaling but also promotes vascular leakage and leukocyte recruitment, precluding its therapeutic utility in treating myocardial ischemia (39), our approach to shift the local balance between flFLT1 and sFLT1 may provide a novel therapeutic approach to not only improve angiogenesis but also constrain inflammation, sensitizing endothelial cells to VEGF signaling without the requirement for detrimental increases in its concentration.

Interestingly, macrophage migration was impaired in *Flt1*^In13pA^ mice, and we also noted a reduction in the VEGF-induced chemotactic responses of *Flt1*^In13pA^ macrophages *in vitro* as well as into an air pouch model *in vivo*. In contrast to ECs, which show only weak FLT1 signaling (18), these data imply cell-type-specific differences in FLT1 responses. It appears that FLT1-mediated signaling, rather than its decoy function, controls the migration of macrophages.

We show that the local balance in flFLT1 and sFLT1 expression represents a critical regulator of angiogenesis and that modulating telescripting to induce a shift in Flt1 isoform expression, could be harnessed to improve angiogenesis, beneficial for tissue repair, e.g., after MI (36). Therapeutically, morpholino phosphorodiamidate oligonucleotides (AMOs) have previously been shown to block U1 binding to the *Flt1* 5’ splice site in intron 13 to generate sFLT1 in cultured cells (3), and could be applied in conjunction with local delivery to the injured tissue by nanobodies or other targeting strategies (40). Intriguingly, by this mechanism, endogenous gene expression is redirected, limiting side effects of unselective overexpression of angiogenic mediators or off-target effects in cells not expressing the transcript (41). Conversely, it would also be interesting in the future to test the effects of rendering the APA signal inactive using AMOs to maintain or enhance flFLT1 expression as a means to restrict the angiogenic response, e.g., in the retina. This may also bypass the reduced viability of mice constitutively lacking sFLT by artificially removing intron 13 to directly connect exons 13-14 of *Flt1* (42). In a broader context, and given the widespread abundance of APA signals (1), our approach to target telescripting at the 5’ splice site to induce APA may serve as a blueprint to further explore the function of truncated transcripts with alternative 3’UTRs. Our study also implies that often overlooked mutations affecting U1 binding to the 5’ splice site could profoundly alter protein functionality, which may be taken into consideration in genetic studies in the future.

Overall, we here demonstrate that telescripting is crucial for the full-length gene expression of FLT1 *in vivo*, and that disruption of this mechanism in the context of hypoxic stress leads to loss of flFLT1, which may serve as a trigger to increase EC responsiveness to angiogenic stimuli during ischemia. In contrast, endogenously generated sFLT1 (in the absence of flFLT1) is sufficient to maintain normal developmental vasculogenesis and pre- and post-natal angiogenesis. Our findings furthermore suggest that modulating telescripting could be used to explore the function of many still uncharacterized truncated gene isoforms generated by APA and indicates that this process is amenable to be harnessed by novel RNA therapeutics to control endogenous gene isoform expression *in vivo*.

## Materials and Methods

### RNA Extraction, Reverse Transcription and qPCR

Total RNA was isolated from different tissues, BMDM and mouse lung endothelial cells using the NucleoSpin RNA kit (Macherey Nagel #740955) or Trizol according to the manufacturer’s instruction. RNA was transcribed using the First Strand cDNA Synthesis Kit (Thermo Scientific, K1612). PCR and qPCR were performed using two reverse primers for *sFlt1* and *flFlt1* (TTAATGTTTGACATGACTTTGTGTG and CGGTTCTTGTTGTATTTTGTGGTTG) and a common forward primer (GCCTTCAATAAAATAGGGACTGTGG). For qPCR, total *Flt1* was amplified using forward primer CACCACGGAGCTCAATACGA and reverse primer CTTCACGCGACAGGTGTAGA. qPCR was performed using a GeneQuant Studio 6 Real-Time PCR System and SybrGreen PCR Mix. Results are expressed as 2^−ΔΔCt^. For absolute qPCR quantification, gBlocks were used as standards. Sequences were as follows:

*s*/*tFlt1*: tgcatgatctacgtgcgtcacatgcagtacGCCTTCAATAAAATAGGGACTGTGGAAAGAAACATAAAATTTTATGTCACAGATGTGCCGAATGGCTTTCACGTTTCCTTGGAAAAGATGCCAGCCGAAGGAGAGGACCTGAAACTGTCCTGTGTGGTCAATAAATTCCTGTACAGAGACATTACCTGGATTCTGCTACGGACAGTTAACAACAGAACCATGCACCATAGTATCAGCAAGCAAAAAATGGCCACCACTCAAGATTACTCCATCACTCTGAACCTTGTCATCAAGAACGTGTCTCTAGAAGACTCGGGCACCTATGCGTGCAGAGCCAGGAACATATACACAGGGGAAGACATCCTTCGGAAGACAGAAGTTCTCGTTAGAGGTGAGCACTGCGGCAAAAAGGCCATTTTCTCTCGGATCTCCAAATTTAAAAGCAGGAGGAATGATTGTACCACACAAAGTCATGTCAAACATTAAaccaatggctttccgagatgCACCACGGAGCTCAATACGAGGGTGCAAATGAGCTGGAATTACCCTGGTAAAGCAACTAAGAGAGCATCTATAAGGCAGCGGATTGACCGGAGCCATTCCCACAACAATGTGTTCCACAGTGTTCTTAAGATCAACAATGTGGAGAGCCGAGACAAGGGGCTCTACACCTGTCGCGTGAAGcactagctcagattcagtagaccgctgttg

*flFlt1*: tgcatgatctacgtgcgtcacatgcagtacGAGCGATGTGGAAGGAGACGAGGATTACAGTGAGATCTCCAAGCAGCCCCTCACCATGGAAGACCTGATTTCCTACAGTTTCCAAGTGGCCAGAGG CATGGAGTTTCTGTCCTCCAGAAAGTGCATTCATCGGGACCTGGCAGCGAGAAACATCCTTTTATCTGAGAACAATGTGGTGAAGATTTGCGACTTTGGCCTGGCCCGGGATATTTATAAGAACCCTGATTATGTGAGGAGAGGAGATACTCGACTTCCCCTAAAATGGATGGCTCCTGAATCCATCTTTGACAAGGTCTACAGCACCAAGAGCGATcactagctcagattcagtagaccgctgttg *flFlt1* was amplified using forward primer GAGCGATGTGGAAGGAGACGAG and reverse primer ATCGCTCTTGGTGCTGTAGACC.

### Morpholino Treatment

Mouse *Flt1* antisense (MoVE) oligonucleotide sequences were 5’-TTTTTGCCGCAGTGCTCACCTCTAA-3’. Oligonucleotides were transfected into 4 × 10^4^ cells using Endoporter (Gene Tools) according to the manufacturer’s instruction to achieve the desired final doses as indicated.

### Generation of Flt1^In13pA^ mice

*Flt1*^In13pA^ mouse line was generated by direct oocytes microinjections using the CRISPR-Cas9 components together with the single-stranded oligo-deoxyribonucleotide (sODN) donor template (FLT1mutTEMPL) with the subsequent chirurgical embryo-transfer. *Flt1*^In13pA^ founder mice were backcrossed into the C57BL/6 background for ten generations.

### Donor DNA template preparation

Donor DNA template FLT1mutTEMPL: (TGCAGAGCCAGGAACATATACACAGGGGAAGACATCCTTCGGAAGACAGAAGTTCTCGTTcGcGGcGAaCACTGCGGCAAAAAGGCCATTTTCTCTCGGATCTCCAAATTTAAAAGCA GG) was chemically synthesized [MWG, eurofins].

### Mouse oocytes microinjections

For the preparation of CRISPR-Cas9 microinjection solution, commercially synthesized FLT1_crR1 with target sequence: CTCGTTAGAGGTGAGCACTG, together with the tracrRNA and, Cas9 protein [Integrated DNA Technologies (IDT), USA] were mixed as follows: 50 pmol of crRNA were mixed with 50 pmol of tracrRNA in 10 mM potassium acetate and 3 mM Hepes (pH 7.5) buffer and incubated at 95^0^C for 1 min, followed by cooling to room temperature. The annealed crRNA/tracrRNA complex was mixed with Cas9 mRNA, Cas9 protein, and FLT1mutTEMPL sODN. The final concentrations of CRISPR-Cas9 components in 0.6 mM Hepes (pH 7.5) and 2 mM potassium acetate microinjection buffer were as follows: crRNA (1 pmol/µl), tracrRNA (1 pmol/µl), Cas9 mRNA (10 ng/µl), Cas9 protein (25 ng/µl), FLT1mutTEMPL sODN donor template (0,04 pmol/µl). The final injection solution was filtered through Millipore centrifugal columns and spun at 20,000*g* for 10 min at +4 °C. Microinjections were performed in B6D2F1 (hybrid between C57BL/6J and DBA strains) in vitro fertilized (IVF) one-cell oocytes. For the IVF oocytes were removed from oviducts of superovulated B6D2F1 female mice in 90 µl of 0,25 mM GSH in mHTF media under the mineral oil. In parallel, *cauda epididymis* and *vas deferens* (*ductus deferens*) of C57BL6 male mouse were transferred to 120 µl of TYH MBCD media and mixed. Ten microliters of sperm solution were transferred to fresh 90 µl of TYH MBCD media and kept for 1 hour at 37°C, 5% CO_2_. Ten microliters of the diluted sperm solution were added to isolated oocytes and placed in incubator at 37°C, 5% CO_2_ for 3 hours. After IVF zygotes were washed 4 times with 120 µl of warm mHTF media and incubated for 3 hours at 37°C, 5% CO_2_. Microinjections were performed in M2 media, using the Transjector 5246 (Eppendorf), and Narishige NT-88NE micromanipulators attached to a Nikon Diaphot 300 inverted microscope. Zygotes that survived microinjections were incubated in KSOM media for overnight at 37°C, 5% CO_2_. Next day two-cell embryos were transferred to oviducts of pseudopregnant CD1 foster mice and carried to term. Positively targeted F0 and F1 animals were identified by qPCR and sequencing analysis of genomic DNA isolated from ear biopsies.

### Flt1^In13pA^ mouse genotyping

TaqMan qPCR analysis was applied for genotyping of *Flt1*^In13pA^ mice. The qPCR reaction was performed using 100 ng of genomic DNA samples, using FLT1_qPCRd1 GGAAGACATCCTTCGGAAGAC, and FLT1_qPCRr1 AATTTGGAGATCCGAGAGAAAA primers in combination with dual-labeled LNA probes 5’-6FAM/3’-IBFQ: (IDT, USA): Flt1wt A<u>G</u>A<u>G</u>G<u>T</u>GA<u>G</u>C (wild type) or Flt1mut CG<u>C</u>GG<u>C</u>GA<u>A</u>C (mutant). The position of LNA nucleotides in probe are underlined. All qPCR reactions were performed in a total volume of 20 μl containing 2 μl of genomic DNA (∼100 ng), 10 μl of 2 X TaqMan Master Mix (Roche), 0,1 μM TaqMan LNA probe and 0,1 μM of each primer. PCR amplification was performed as follows: enzyme activation at 95°C for 5min (Ramp Rate (RR) 4,4), with subsequent 50 cycles of qPCR: 95°C – 10sec (RR 4,4); 60°C – 30sec (RR 2,2) and final cooling cycle: 40°C – 10sec (RR 2,2). PCR products from qPCR positive *Flt1*^In13pA^ mice were confirmed by sequencing.

### Hind limb ischemia

12 week old *Flt1^+/+^* and *Flt1^In13pA^*mice were anesthetized with isoflurane (2%) and placed on 37°C heated pads during surgery. Hair was removed from the hindquarters and the femoral artery was exposed and isolated from the femoral vein and nerve. Unilateral hindlimb ischemia was induced by partial excision of left femoral artery. After 14 days mice were sacrificed and hind limb muscles were excised for further analysis. All animal studies and numbers of animals used conform to the Directive 2010/63/EU of the European Parliament and have been approved by the appropriate local authorities (Regierung von Unterfranken, Würzburg, Germany, Akt.-Z 55.2.2-2532-2-1272).

### Myocardial infarction

Myocardial infarction was induced in 8 to 12 week old male C57BL/6 mice by permanent ligation of the left descendent coronary artery. Mice received buprenorphine, 0.1mg/kg i.p. as analgesia, while anesthesia was induced by inhalation of isoflurane (4.0%). Mice under deep anesthesia were intubated with an endotracheal cannula and placed under mechanical ventilation (CWE INC. SAR-830-AP; 130 respirations/min, peak pressure max. 18 cm H2O) on a heating pad to maintain body temperature. 1.5-2.5% isoflurane was used to maintain anesthesia during surgery. The heart was exposed via a 4th intercostal thoracotomy. The left descendent coronary artery was visualized and ligated with a 7/0 non-resorbable nylon suture. The thorax was closed with 3 separated 6/0 non-resorbable nylon sutures and the skin with a continuous 6/0 non-resorbable nylon suture. Buprenorphin injections were administered i.p. 6 hours after the surgery and twice daily in the two days following surgery. Exclusion criteria related to general animal health had been defined upon pre-registration of the experiments for approval by local authorities. Briefly, animal well-being was controlled at regular intervals. The well-being of the animals was monitored with a cumulative point system (Score Sheet), with a pre-defined threshold associated with early termination of experiments and humane euthanasia of the animals. For myocardial infarction studies, mice dying after myocardial infarction before the planned sacrifice date were excluded from analysis after 28 days. At sacrifice, specific exclusion criteria related to the size of the infarct were applied: mice with the absence of a visible infarct or mice showing small infarcts (below∼30% of the left ventricle, as macroscopically determined by the experimentator after sacrifice) were excluded from analysis; applying these criteria, we excluded two mice from the study.

For the analysis of cardiac function, anesthetized mice were examined using a Vevo2100 (VisualSonics) high resolution imaging device and a 30-MHz probe (43). Briefly, animals were monitored by surface ECG using limb electrodes and body temperature (37 °C warming plate, anal probe). Mice were shaved and fixed on the warmed ECG plate in prone position. B-mode movies of the parasternal long axis (PSLAX) and a orthogonal short axis at the proximal level of the papillary muscle were acquired. For the analysis of end-diastolic and end-systolic volumes, the endocardium of the left ventricle was traced at both diastole and systole. An integrated software tool (Simpson) was used for analysis of ventricular volumes and FAC. EF was calculated using the formula EF = ((EDV − ESV)/EDV) × 100, where EDV is end diastolic volume. Mid-ventricular M-mode was used to determine the systolic (s) and diastolic (d) left ventricular internal diameter (LVID) and to calculate fractional shortening (FS %). Mice with heart rates below 430 beats per minute were excluded from the analyses.

Mice were killed by cervical dislocation under isoflurane anesthesia. All animal studies and numbers of animals used conform to the Directive 2010/63/EU of the European Parliament and have been approved by the appropriate local authorities (Regierung von Unterfranken, Würzburg, Germany, Akt.-Z 55.2.2-2532-2-1272).

### ELISA

4×10^4^ mouse lung endothelial cells isolated from *Flt1^+/+^* and *Flt1^In13pA^*mice were plated on 24 well plates in complete EC media. After 24h cells, when cells reached confluence, medium was changed to 1% FCS media. Cell were incubated for an additional 28 h, supernatant was collected, centrifuged at 400 g for 5 min, and sFLT1 concentrations were measured by ELISA as specific by the manufacturer (R&D, #DY471).

### 3’ RACE Experiments

3’ RACE reactions were performed on RNA extracted from mouse liver tissue and the RACE experiments were done according to the manufacturer’s instructions (3’ RACE System for Rapid Amplification of cDNA Ends, Invitrogen).

### Mouse lung endothelial cell isolation

Mouse lung endothelial cells were isolated from mouse lungs of 6-8 week old mice. Three milliliters of cold PBS was injected via the heart to flush the lung of blood cells. Lungs were removed, cut in small pieces and incubated with dispase II (25U) for 1h at 37°C, with vigorous shaking at 350 rpm. Cell suspension was filtered through 70 µm strainer and washed with 50 ml of DMEM-F12 with 10% FCS, 100 U/mL penicillin and 100 μg/mL streptomycin. Cells were pelleted by centrifugation for 5 min at 300 g and plated in gelatin-coated flasks in complete EC media – DMEM-F12 with 20% FCS, 100 U/mL penicillin and 100 μg/mL streptomycin with addition of ECGF (12 µg/ml, PeloBiotech #PB-ECGF-1 Heparin). After reaching confluence, cells were detached by accutase (Sigma Aldrich #A6964) and endothelial cells were selected using CD31 murine MicroBeads according to the manufacturer instructions (Miltenyi, #130-097-418). Cells were further cultivated on gelatin-coated plates in complete EC media.

### Tube Formation assay

48 well plates were covered with 50 µl of Geltrex previously thawed overnight at 4°C (Gibco # A1413201) and incubated for 30 minutes at 37°C. 3×10^4^ mouse lung endothelial cells isolated from *Flt1^+/+^*and *Flt1^In13pA^* mice were plated per well in complete EC media and incubated for 24 h at 37°C and 5% CO_2_. Images were taken by a Nikon eclipse Ts2 microscope and number of meshes were counted per microscopic field. To investigate the effect of exogenously added VEGF, 3×10^4^ mouse lung endothelial cells isolated from *Flt1^+/+^* and *Flt1^In13pA^* mice were plated in Geltrex-coated 48 well plates in supernatant collected from *Flt1^+/+^* and *Flt1^In13pA^* mouse lung endothelial cell cultures containing sFLT1 (*Flt1^+/+^* supernatant: 2000 pg/ml sFlt1; *Flt1^In13pA^* 5242 pg/ml sFlt1) in DMEM-F12 with 5% FCS, 100 U/mL penicillin and 100 μg/mL streptomycin. Cells were and treated with 50 ng/ml of recombinant murine VEGF (Peprotech #450-32). After 24 h of incubation, images were recorded by a Nikon eclipse Ts2 microscope. To investigate if sFlt1 produced by *Flt1^In13pA^* EC is able to repress VEGF-induced angiogenesis, 2×10^4^ SVEC cells were plated on Geltrex-coated 48 well plates in DMEM-F12 (0.5% FCS) or in supernatant collected from *Flt1^+/+^* and *Flt1^In13pA^* mouse lung endothelial cell cultures with addition of 50 ng/ml of recombinant murine VEGF. Cells were incubated for 24 h at 37°C and 5% CO_2_. Images were taken after 2, 4, 6 and 24 h by a Nikon eclipse Ts2 microscope. The number of tubes was counted per microscopic field.

### Endothelial cell proliferation

2×10^3^ mouse lung endothelial cells were plated in gelatin-coated 48-well plates in complete EC media and incubated for 48 h at 37°C and 5% CO2. Cells were then washed with PBS, detached by accutase, centrifuged and resuspended in 20µl of PBS. 10µl of cell suspension was diluted in 90µl of Tryptan Blue solution (Sigma #T8154) and numbers of live cells were counted using a hemocytometer.

### In vitro macrophage migration assay

Bone marrow-derived macrophages were flushed from mouse femur and tibia, filtered through a 70 µm nylon mesh and plated in 10 cm dishes at a density of 2×10^6^/ml in 10ml. Cells were differentiated into macrophages in medium containing RPMI1640, 15% L929-conditioned medium, 10% fetal bovine serum (heat inactivated at 56 °C/45min), 50 µM β-mercaptoethanol (Gibco #31350-010), 100 U/mL penicillin, and 100 μg/mL streptomycin for 7 days. Before migration assay cells were starved for 4 hours in RPMI medium containing 0,5% FCS. Accutase was used to harvest the cells and 10^5^ cells were transferred to the transwell inserts (100µl of cell suspension in 0.5% FCS medium). Cells were allowed to settle down for 30min before adding attractants to the receiver well - 50ng/ml VEGF (Peprotech #450-32) or 100ng/ml CCL2 (Peprotech #250-10). Cells were incubated 4h at 37 °C and 5% CO_2_. After migration, 4% PFA was used to fix the cells and they were stained with 1% crystal violet. Transwell inserts were washed in distilled water to remove excess crystal violet and non-migrated cells on top of the membrane were removed with cotton swabs. Images were taken by Leica DM4000 microscope. Macrophage migration index was calculated as a number of cells migrated toward attractant divided by the number of spontaneously migrated macrophages in negative control.

### Air pouch model

Mice were anesthetized with isofluran and 5 ml of sterile air was subcutaneously injected in the intrascapular area of the back. On day 3, an additional 3 ml of sterile air was injected. On day 5 sterile PBS or PBS + 50 ng/ml VEGF (Peprotech #450-32) was injected into the air pouch and 24 h later exudates were collected. The number of leukocytes in the exudate was counted using a Neubauer counting chamber and a flow cytometric analysis was performed.

### Immunostainings of Retinal Endothelial Cells

Eyes were fixed in 4% PFA for 15 minutes. The retinal cups were dissected and fixed for an additional 2 hours at 4°C. Retinal EC were then stained overnight with FITC-conjugated isolectin B4 from Bandeirae simplicifolia (0.04 mg/ml; Sigma-Aldrich #L9381) in Pblec Solution (1 mM MgCl2, 1 mM CaCl2, 0.1 mM MnCl2, 1% Triton X-100 in PBS) at 4°C. The retinal cups were washed in PBS, cut into quadrants and flat-mounted on microscopic slides with Mowiol solution (Dianova). Images were taken by fluorescent microscope (Olympus, Hamburg, Germany). To study physiological vascular development at P5 and P7, the vascular and entire retinal areas of each quadrant were analysed using ImageJ and Angiotool software.

### 3’mRNA Sequencing and Analysis

The total RNA from mouse lung endothelial cells was analyzed by 3’mRNA sequencing with QuantSeq 3’ mRNA-Seq Library Prep Kit REV for Illumina (Lexogen #016.24). Library preparation and sequencing was done according to the manufacturer’s instruction. Samples were sequenced in triplicates. FASTQ files went through pre-processing according to Lexogen’s indications. Briefly, reads were trimmed with Cutadapt(44) to remove residual technical sequences. The trimmed reads were then aligned against the mouse genome (Gencode M29) with STAR(45). PolyA site read counts for Flt1 were obtained using the Samtools(46) view command with a filter for mapping quality lower than 30 applied. Genomic coordinates were used to determine the intervals where reads were counted. These were obtained from the PolyASite(47) database for known sites and determined as a 10 bps interval around putative unannotated polyA sites. PolyA sites were examined by observing the sequence composition and read coverage with the IGV(48) genome viewer. Proportions (in percentages) were then calculated on the read counts sum for all valid isoforms. RNA sequencing data were analyzed by R software including principal component analysis, DESeq2 analysis, and Gene Set Enrichment Analysis (GSEA). Downstream statistical analysis of differential gene expression was conducted using the DESeq2 Bioconductor package (p adjusted ≤ 0.001, log2cutoff ≤ 0.5). Angiogenesis-related genes presented in the heatmap were obtained from the molecular signature database (mouse gene set - Hallmark_Angiogenesis, Gene Set Enrichment Analysis, www.gsea-msigdb.org) and the Gene Ontology Term angiogenesis (GO:0001525).

### Flow cytometry

For air pouch analyses, exudates were collected and centrifuged. Cells were stained for surface markers with specific antibodies: anti-CD45 (Biolegend #103128, dilution 1:400); anti-CD11b (Biolegend #101237, dilution 1:400), anti-CD115 (Invitrogen #17115280, dilution 1:300), anti-Ly6G (BD #560603, dlution 1:400), anti-Ly6C (Biolegend #128033, dilution 1:400), anti-CCR2 (Biolegend #150609, dilution 1:20), anti-F4/80 (Biolegend #123114, dilution 1:100), anti-CD64 (Biolegend #139316, dilution 1:100), anti-MHCII (Biolegend #BV785, dilution 1:100). Probes were analyzed using a FACSCelesta and FlowJo 10.0 software (BD Biosciences).

Endothelial cell purity was analyzed by staining of surface markers anti-CD31 (Invitrogen #11-0311-82, dilution 1:300) and anti-CD144 (Miltenyi #130-128-207, dilution 1:10). 3 days after MI, hearts were removed and enzymatically digested in RPMI medium containing 450 U/ml collagenase I (Sigma-Aldrich #C0130), 125 U/ml collagenase XI (Sigma-Aldrich #C7657), 60 U/ml Hyaluronidase (Sigma-Aldrich #H3506) for 1 hour at 37°C with agitation. Thereafter digested material was passed through a 70 µm cell strainer. Unspecific binding to Fc receptors was blocked by incubation of cell suspension with anti-CD16/32 (Biolegend #101302, dilution 1:50) at 4°C for 10 minutes. After pelleting, cells were stained with Fixable Viability Dye (ThermoFisher, 1:1000) and fluorochrome conjugated antibodies against anti-CD45 FITC (Invitrogen # 11-0451-82, dilution 1:300), anti-CD11b PercpCy5.5 (BioLegend #101228, dilution 1:400), anti-Ly6C BV510 (BioLegend #128033, dilution 1:300), anti-Ly6G Pacific Blue (BioLegend #127612, dilution 1:100), CD115-APC (Invitrogen #17-1152-82, dilution 1:300), anti-F4/80 PeCy7 (Biolegend #123114, dilution 1:300), anti-TIM4 (BD Bioscience #564147, dilution 1:100) and anti-MHCII BV785 (Biolegend, #107645, dilution 1:300) at 4°C for 30 minute. Results were acquired using a FACS Celesta (BD Biosciences) and analyzed with FlowJo 10.5.3.

### Histology and immunohistochemistry

Hind limb muscles and hearts were fixed overnight in 4% para-formaldehyde and embedded in paraffin. 5 µm sections were used for immunohistochemistry. Paraffin was removed from sections by consecutive washes with xylene and rehydratation in decreasing ethanol %. Citrate antigen retrieval and 1 h of blocking with anti-mouse CD16/32 (Biolegend #101302, dilution 1:50) was performed before staining. Endothelial cells were stained with anti-CD31 antibody (R&D #AF3628, dilution 1:70) and macrophages with Mac2 antibody (Cederlane #CL8942AP, dilution 1:500). The following day, slides were washed in PBS 3×5 minutes and then stained with the secondary antibody Goat anti-rabbit AlexaFluor555 (Invitrogen #A21429, working concentration 2 µg/µl) or Goat anti-rat Alexa488 (Thermofisher #A-11006, working concentration 4 µg/µl) in PBS for 1h. To visualize cardiomyocytes, Rhodamine (Vector Lab. # RL-1022) was added together with secondary antibody. After washing in PBS (3×5 min) nuclei were counterstained with DAPI (Vectashield #H-1200). Images were taken using a Leica DM4000 microscope and analyzed using Diskus software.

### Protein lysate preparation and Western blotting

8×10^4^ mouse lung endothelial cells isolated from *Flt1^+/+^* and *Flt1^In13pA^*mice were plated on gelatin-covered 12-well plates. After 24 h, cells were washed 3 times with ice-cold PBS and then lysed in ice-cold RIPA buffer (20 mM Tris pH 7.4, 150 mM NaCl, 1 % Na-Deoxycholat, 1 % TritonX 100, 0.1 % SDS, 10 mM EDTA, 0.5 mM Na-Orthovanadat, 10 mM Na-Pyrophosphat, 0.5 mM p-Nitrophenylphosphat). Samples were vortexed and kept on ice for 30 min. Thereafter we proceeded with sonication for 1 minute at a power of 180 W (10 seconds sonication/10 seconds rest for each cycle). Samples were centrifuged at 10,000 x g for 20 minutes at 4°C to pellet cell debris, and supernatant was transferred to a fresh microfuge tube. Total protein concentration was determined by BCA kit (Thermo Fisher #23225). For Western blotting, lysates were mixed with 2X sodium dodecyl sulphate (SDS) sample buffer to a final dilution of 1X and heated at 95°C for 5 minutes. 40 µg of protein lysate was separated by 8% SDS polyacrylamide gel electrophoresis following standard protocols and transferred to nitrocellulose membranes. Membranes were blocked with 3% BSA in Tris-buffered saline with 0.1% Tween 20 (TBST) for 1 h followed by overnight incubation at 4°C with primary antibodies: anti-Vegfr1 (Bioss #bs-0170R, dilution 1:500), anti-phosphoY1175-Vegfr2 (Biorbyt #0rb256644, dilution 1:500), anti-Vegfr2 (Cell Signaling #2479, dilution 1:1000) and anti-actin (Santa Cruz #sc-47778, dilution 1:1000). Membranes were then washed three times in TBST and probed with horseradish peroxidase-conjugated goat anti-rabbit or goat anti-mouse antibody (Biorad, Feldkirchen, Germany). Vizualisation was done using enhanced chemiluminescence reagent (GE Healthcare, München, Germany) and chemiluminescence images were taken using the Amersham Imager 600 (GE Healthcare, München, Germany).

### Statistics

Results are presented as mean ± SD, and were analyzed using Prism9 Software (GraphPad, Inc, La Jolla, CA). The unpaired Student’s t-test was used for 2-group comparisons with normally distributed data. For multiple group comparisons of normally distributed data, the 2-way ANOVA followed by the multiple-comparison Tukey’s post hoc test was used. p<0.05 was considered statistically significant. The individual sample sizes for each set of data is provided in the figures and their legends.

### Data availability

The data that support the findings of this study are available from the corresponding author upon request. Raw bulk sequencing data will be deposited to Gene Expression Omnibus repository.

## Author contributions

M.B. and S.V. designed the study, acquired and analyzed data, and wrote the manuscript; J.U., L.R., L.H., M.E., J.B., S.S., N.A.S., and A.R., acquired and analyzed data; G.R., C.C., W.K., M.W., M.K., E.H. and K.L. acquired and analyzed data and critically revised the manuscript; B.V.S. and T.S.R. generated the mice; R.B. processed sequencing data; A.Z. designed the study, analyzed data, and wrote the manuscript.

## Acknowledgments

We thank Yvonne Kerstan, Petra Hönig-Liedl and Melanie Rösch for expert technical assistance, and Ebba Brakenhielm for critical review of the manuscript. This work was financially supported by the Interdisciplinary Center for Clinical Research (Interdisziplinäres Zentrum für Klinische Forschung (IZKF), University Hospital Würzburg (E-352 to S.V., and A.Z.), the Alexander von Humboldt-Stiftung (Humboldt Research Fellowship to M.B.), and the Deutsche Forschungsgemeinschaft (DFG, German Research Foundation, projects 453989101-CRC1525 to C.C., W.K., K.L., M.K., and A.Z., DFG KU 1037/13-1 to M.K., 324392634-TR221, 374031971-TRR 240, ZE827/14-1, #396923792, #432915089, and #505700170 to A.Z).

