## Supplemental Figures for "Redirecting full-length FLT1 expression towards its soluble isoform promotes postischemic angiogenesis"

**Supplemental Figure 1**

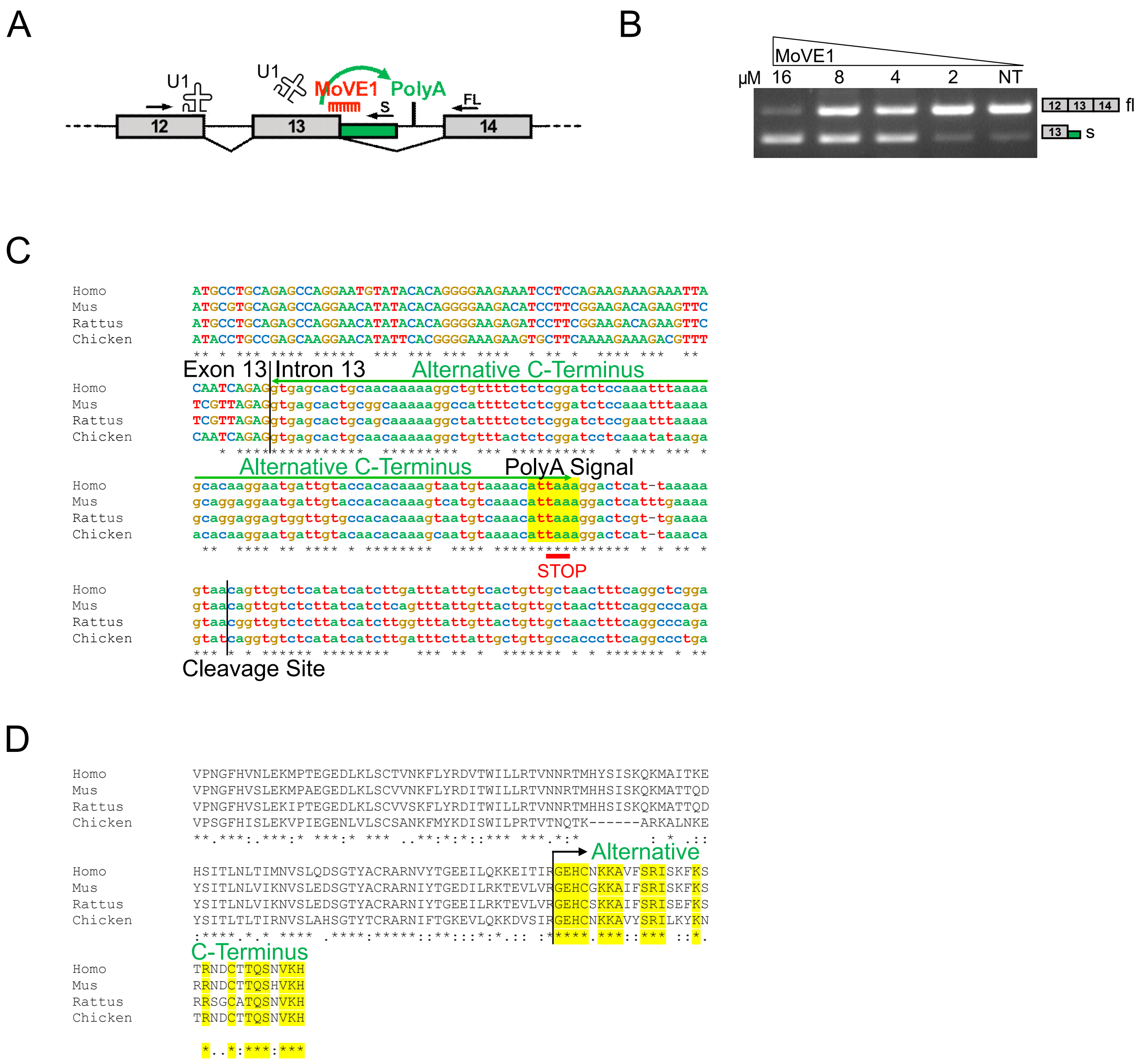


**Supplemental Figure 1**. (**A**) Schematic representation of morpholino oligonucleotide (MoVE1) binding to *Flt1* intron 13/exon13 5′-splice site thereby blocking binding of U1 snRNP resulting in activation of an alternative cleavage-polyadenylation (polyA) signal within intron 13. (**B**) Dose-dependent treatment of mouse SVEC with MoVE1results in increased *sFlt1* and concomitant decreased *flFlt1* transcript expression compared to non-treated (NT) cells. (**C**) Interspecies conservation of *Flt1* intron 13 nucleotide sequence and (**D**) of C terminal sFLT1 protein.

### Supplemental Figure 2

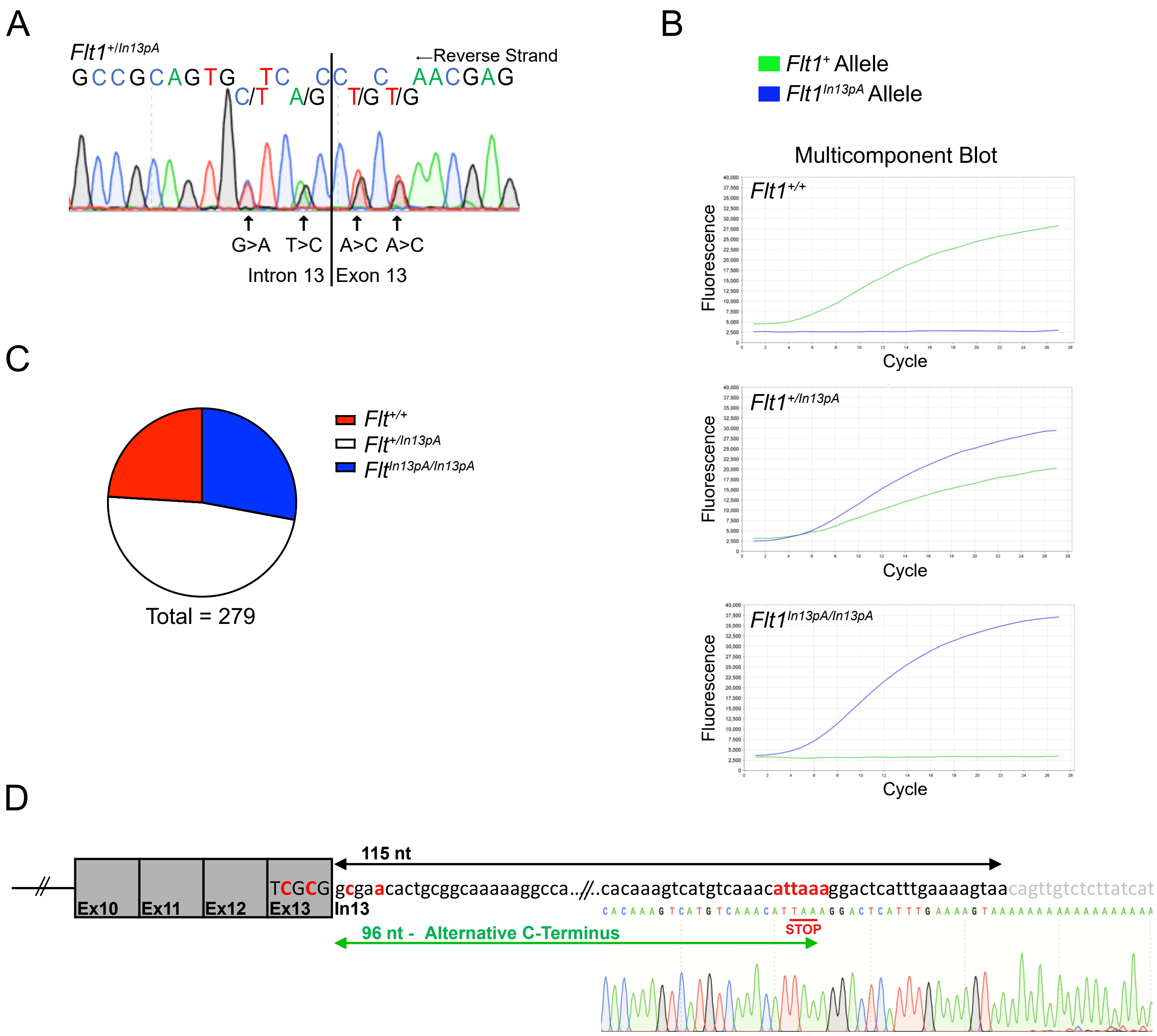


**Supplemental Figure 2.** (**A**) Genomic DNA sequencing chromatogram of heterozygous *Flt1*$^{+/In13pA}$ F1 founder mice. (**B**) TaqMan multicomponent plot obtained from genotyping *Flt1*$^{+/In13p}$ mice carrying one *Flt1*$^{+}$ and *Flt1*$^{In13p}$ allele. **(C)** Genotype distribution of offspring from heterozygous (Flt1$^{+/In13pA}$) parental mice. **(D)** Schematic representation and sequencing results of the oligo(dT) primed RACE-PCR product from liver tissue cDNA, showing a read through of 115 nucleotides in intron 13, encoding an alternative 32 amino acid sFlt1 tail, demonstrating correct alternative pre-mRNA processing in Flt1In13pA mice. Flt1In13pA genomic mutations and the putative polyadenylation signal hexamer are depicted in red.

#### Supplemental Figure 3

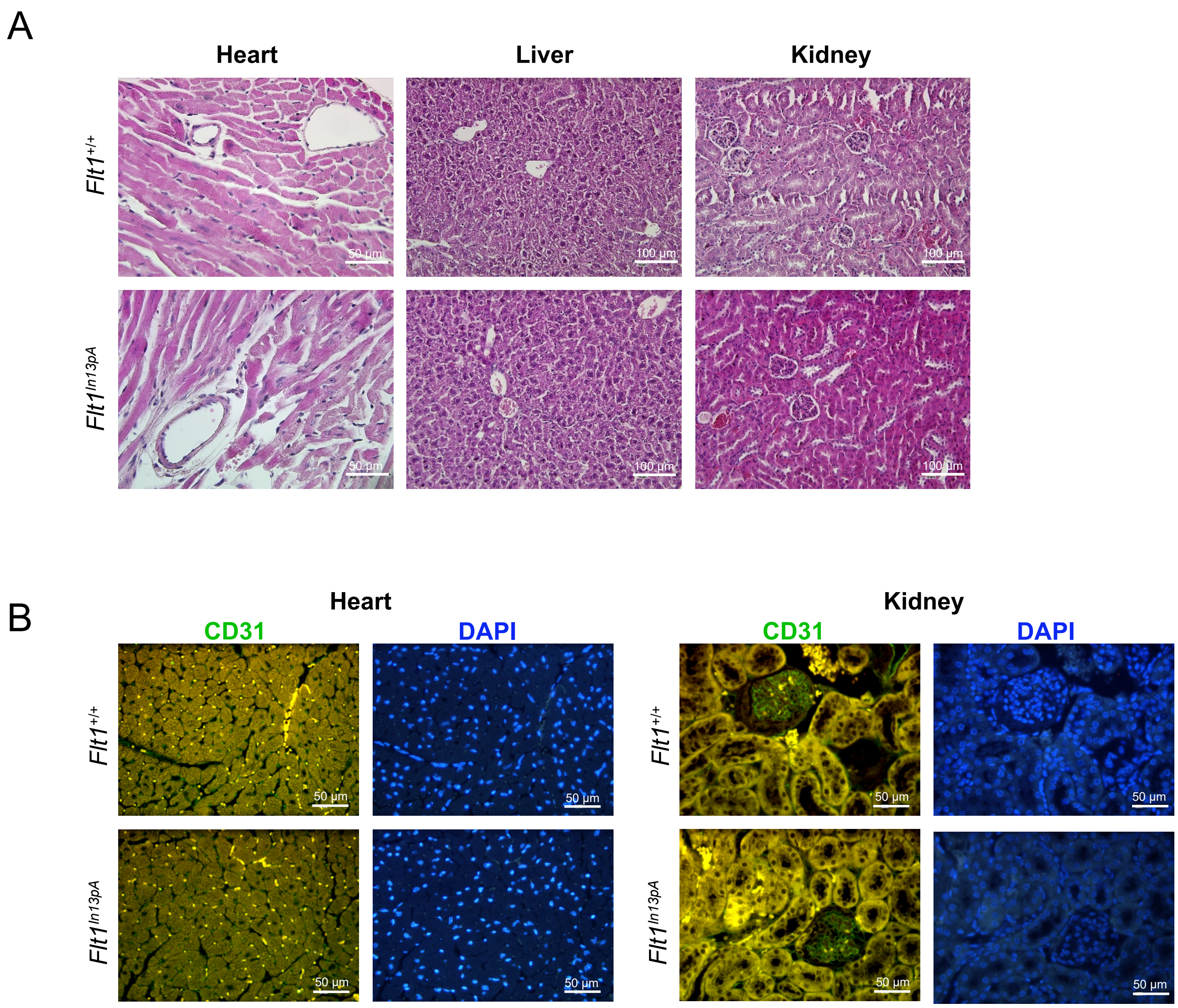


**Supplemental Figure 3.** (**A**) Tissues sections of Flt1$^{+/+}$ and Flt1$^{In13pA}$ heart, liver and kidney stained with H&E; scale bars for heart, 50µm, and for liver and kidney, 100µm. (**B**) Representative immunofluorescence staining of CD31 and DAPI in heart and kidney tissue sections from Flt1$^{+/+}$ and Flt1$^{In}$13pA mice (scale bars, 50 µm).

### Supplemental Figure 4

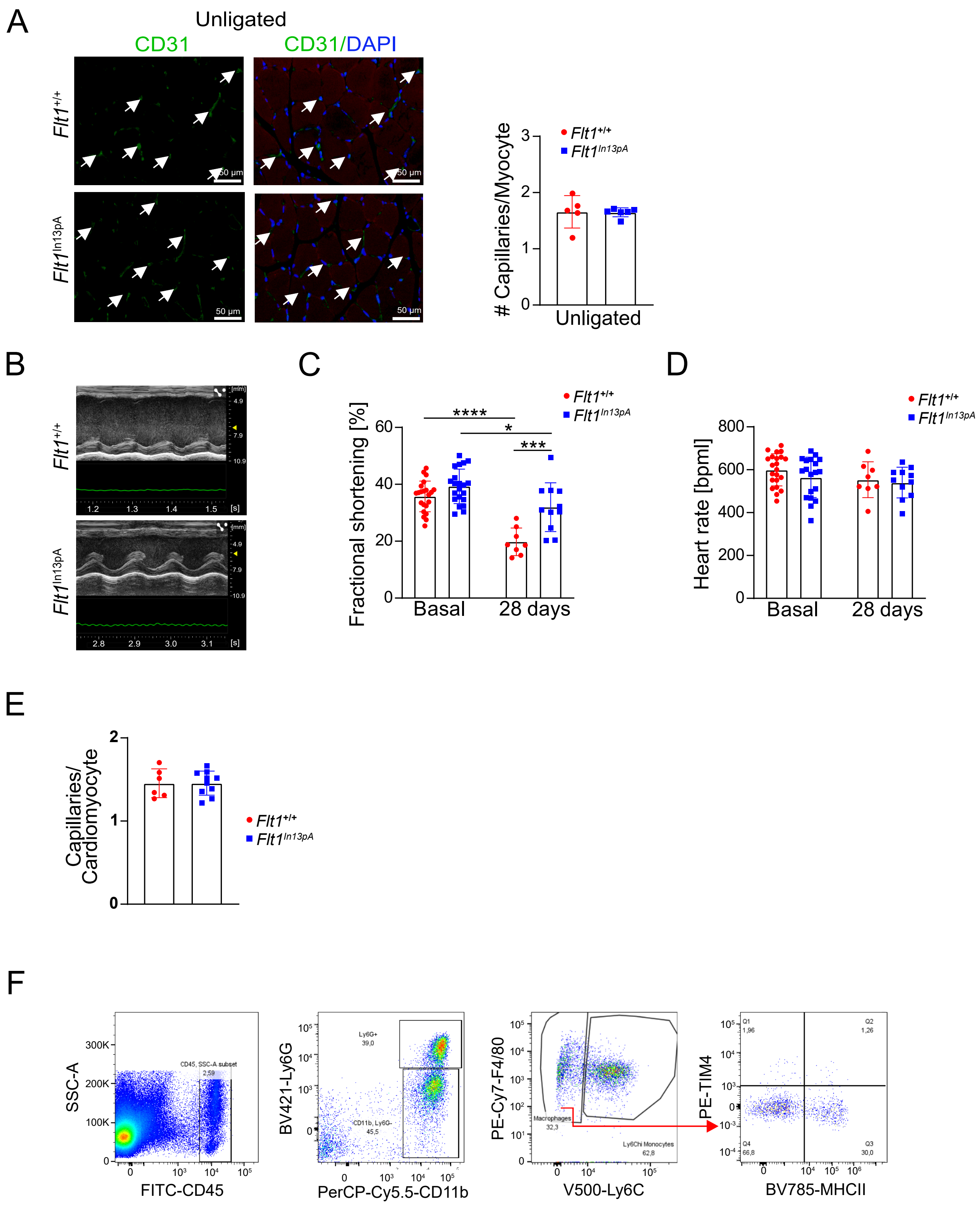


**Supplementary Figure 4.** (**A**) Representative immunofluorescence staining and quantification of CD31$^+$ capillaries in unligated gastrocnemius muscle. Results were analyzed by two-tailed, unpaired Student t-test. (B-F) Flt1$^{+/+}$ and Flt1$^{In13pA}$ mice were subjected to MI injury. (**B**) M-Mode echocardiogram and (**C**) quantification of fractional shortening (%) before (n=22 Flt1$^{+/+}$ and n=21 Flt1$^{In13pA}$ mice) and at 28 days post-MI (n=8 Flt1$^{+/+}$ and n=11 Flt1$^{In13pA}$ mice). Results were analyzed by One way ANOVA followed by Tukey's post-hoc test. (**D**) Heart rate before (n=22 Flt1$^{+/+}$ and n=19 Flt1$^{In13pA}$ mice) and at day 28 post-MI (n=8 Flt1$^{+/+}$ and n=11 Flt1$^{In13pA}$ mice). Data were analyzed by One way ANOVA followed by Tukey's post-hoc test. bpm – beats per minute. (**E**) Quantification of CD31$^+$ capillaries relative to the number of cardiomyocytes in the remote myocardium (n=6 Flt1$^{+/+}$ and n=10 Flt1$^{In13pA}$ mice) at day 28 after MI. Results were analyzed by two-tailed, unpaired Student t test (**F**) Gating strategy to evaluate monocyte, macrophage and neutrophil counts in the heart 3 days post MI. Data represent mean ± SD. *p<0.05, **p<0.01, ***p<0.001, ****p<0.0001.

**Supplemental Figure 5**

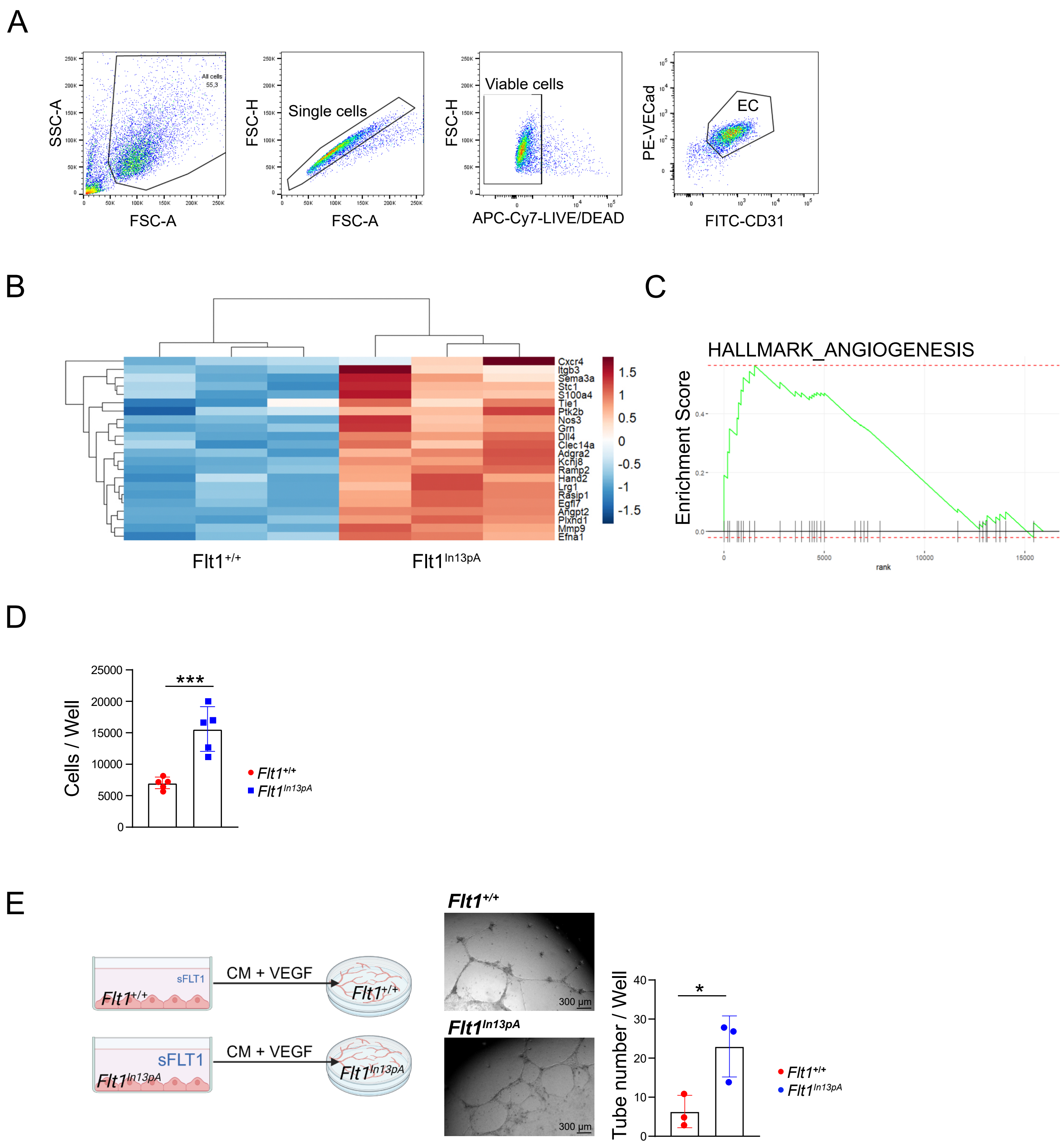


**Supplemental Figure 5.** (**A**) Gating strategy to evaluate the % of CD31+CD144+ primary mouse lung endothelial cells amongst total live cells by flow cytometry. (B-C) Bulk RNA-sequencing of $Flt1^{+/+}$ and $Flt1^{In13pA}$ lung endothelial cells exposed to hypoxia over 24 h (n=3 experiments performed with pooled endothelial cells isolated from n=8 mice) (**B**) Heatmap of RNA-seq data for angiogenesis-associated genes. (**C**) Gene set enrichment analysis (GSEA) plot of angiogenesis gene set. (**D**) Endothelial cells proliferation. $Flt1^{+/+}$ and $Flt1^{In13pA}$ mouse lung endothelial cells were grown for 48 h, detached and counted (independent endothelial cell isolations from n = 5 mice per genotype). Results were analyzed by two-tailed, unpaired Student t test. (**E**) Tube formation assay of $Flt1^{+/+}$ and $Flt1^{In13pA}$ mouse lung endothelial cells incubated with VEGF supplemented
